# Dissecting the subtropical adaptation traits and cuticle synthesis pathways via the genome of the subtropical blueberry *Vaccinium darrowii*

**DOI:** 10.1101/2021.09.03.458838

**Authors:** Fuqiang Cui, Xiaoxue Ye, Xiaoxiao Li, Yifan Yang, Zhubing Hu, Kirk Overmyer, Mikael Brosché, Hong Yu, Jarkko Salojärvi

## Abstract

*Vaccinium darrowii* is a subtropical wild blueberry species, which was used to breed economically important southern highbush cultivars. The adaptation traits of *V. darrowii* to subtropical climate would provide valuable information for breeding blueberry and perhaps other plants, especially against the background of global warming. Here, we assembled the *V. darrowii* genome into 12 pseudochoromosomes using Oxford Nanopore long reads complemented with Hi-C scaffolding technologies, and predicted 41 815 genes using RNAseq evidence. Syntenic analysis across three *Vaccinium* species revealed a highly conserved genome structure, with the highest collinearity between *V. darrowii* and *V. corymbosum*. This conserved genome structure may explain the high fertilization during crossbreeding between *V. darrowii* and other blueberry cultivars. Gene expansion and tandem duplication analysis indicated possible roles of defense and flowering associated genes in adaptation of *V. darrowii* to the subtropics. The possible *SOC1* genes in *V. darrowii* were identified with phylogeny and expression analysis. Blueberries are covered in a thick cuticle layer and contain anthocyanins, which confer their powdery blue color. Using RNA-sequencing, the cuticle biosynthesis pathways of *Vaccinium* species were delineated here in *V. darrowii*. This result could serve as a reference for breeding berries with customer-desired colors. The *V. darrowii* reference genome, together with the unique traits of this species, including diploid genome, short vegetative phase, and high compatibility in hybridization with other blueberries, make *V. darrowii* a potential research model for blueberry species.

## Introduction

Blueberry is rich in bioactive compounds, such as anthocyanins, flavonoids, and other phenolic compounds^1,2^. Diets rich with blueberries have shown multiple health-promoting properties, including antitumor, antibacterial, anti-inflammatory effects, as well as neural, cardiovascular and retinal protection^1,2^. Together with their taste, these characteristics have promoted blueberry consumption and thereby steadily increased market demand. World blueberry production has more than doubled from 300 000 tons in 2008 to 657 000 tons in 2017 ^3^. In China, production has increased more than 100-fold between 2006 and 2015, and reached 234 700 tons in 2020^4,5^. Two major cultivar categories among blueberry crop species are northern highbush and southern highbush, which constitute the majority of blueberry production worldwide^5–7^. Southern highbush blueberry breeding has made considerable progress over recent years both in terms of newly-bred cultivars and growth area^5^. Notably, the southern highbush cultivars have a lower chilling requirement for flower induction, making it possible to produce berries year-round with improved management techniques^8,9^. Thereby, the southern highbush blueberry is a highly attractive emerging crop species for both farmers and researchers.

*Vaccinium darrowii* is an evergreen shrub originating from the subtropical area of America^10^. The crossbreeding between *V. darrowii* and the northern highbush blueberry (*V. corymbosum*) generated the southern highbush cultivars, which have expanded the blueberry-producing region to subtropical/tropical areas^10,11^. In addition to the practical use of *V. darrowii* in breeding, it is a good model for the study of plant adaptation to subtropical climates. Additionally, *V. darrowii* has multiple advantages for basic research, such as a short vegetative phase, with seedlings that usually flower within one year in the wild. This species is self-compatible in fertilization: berries develop from up to 20% of flowers after self-pollination, and 70% in crosspollination^12^, and self-pollinated berries produce 21 plump seeds on average^12^. In contrast to the tetraploid northern highbush, *V. darrowii* is mostly diploid (2n = 2x = 24) in nature^13^. These properties make *V. darrowii* an ideal candidate genetic model for the *Vaccinium* genus, especially for species growing in subtropical regions.

Taxonomically, blueberries make up the *Cyanococcus* section under the genus *Vaccinium*. The northern highbush blueberry, *V. corymbosum* (2n = 4x = 48), is the only species in the *Cyanococcus* section with a chromosome-level genome assembly^14^. Species in other sections of *Vaccinium*, such as cranberry (*V. macrocarpon*), a species sharing similar geographical distribution with the *V. corymbosum* in the temperate region, and the boreal bilberry (*V. mytillus*) have recently been sequenced^15,16^. These genomes serve as valuable resources for genome-enabled breeding of northern *Vaccinium* species, while genome information for southern adapted blueberries is still lacking. Here we sequenced the genome of wild *V. darrowii*. A high quality genome assembly will assist studies on southern blueberry cultivars, which have a shorter vegetative phase and lower chilling requirement compared to northern cultivars^17,18^. Another advantage of *V. darrowii* is its compatiblity for hybridization with other blueberry cultivars regardless of their ploidy levels^12,19^. This enables easy transfer of desired genetic traits from V. *darrowii* to any targeted cultivar.

## Results

### Genome sequencing and assembly

*Vaccinium darrowii* is an evergreen dwarf-shrub in the subtropics^10^. It is early flowering, usually flowering in the spring of the second year after planting (Fig. 1A). Both berries and leaves are covered by apparent waxy layers (Fig. 1B and C). Multiple studies have shown the chromosome number and genome size of *V*. *darrowii* to be around 550 Mb with 12 chromosomes^19,68–70^. Consistently, our flow cytometry analysis estimated the genome size of *V. darrowii* to be ~ 560.5 Mb (Fig. S1). A K-mer based analysis with Illumina short read sequencing yielded 555.38 Mb as the estimated genome size with 42.92% repeats. The heterozygosity rate was 1.27%, which is a common level of heterozygosity for wild woody species (Fig. S2).

**Fig. 1.**
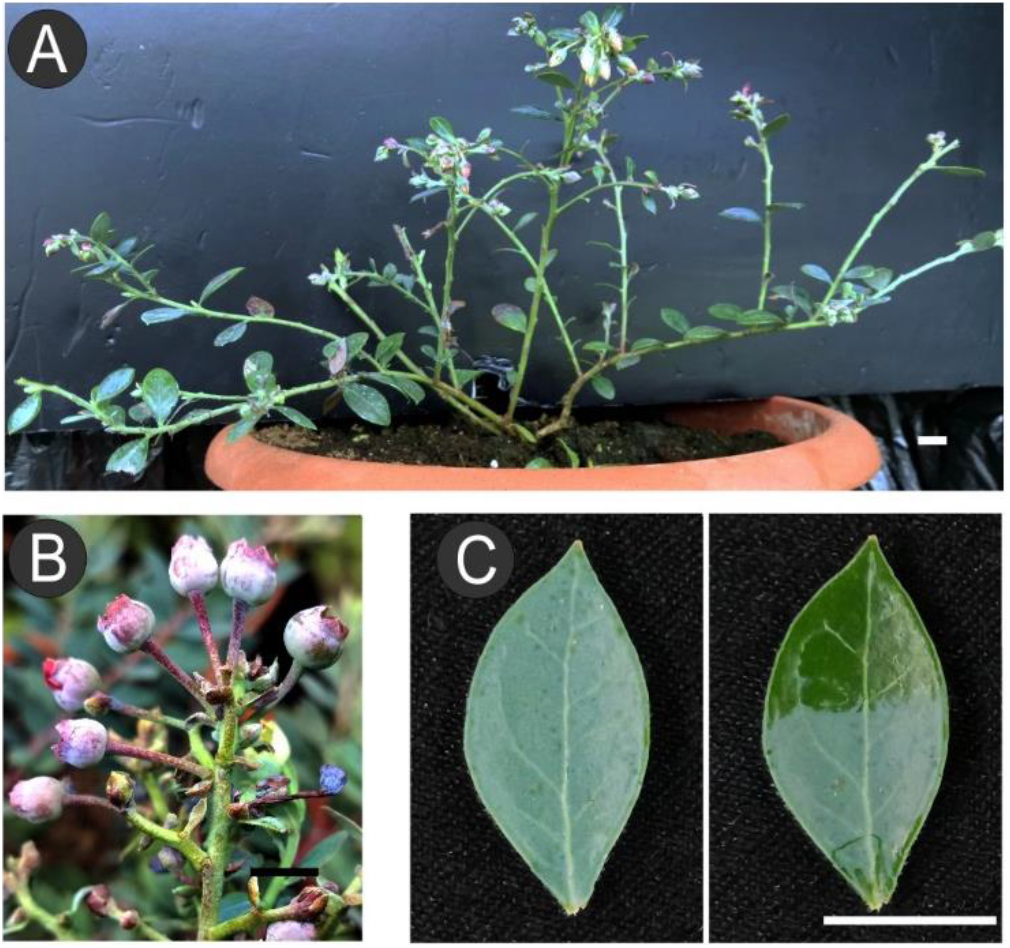
*V. darrowii* phenotypes. (A) Cutting propagated plants flowered at 14-months of age. (B) Waxy berries set on the pedicels. (C) A typical waxy leaf of *V. darrowii*. Intact leaf (left); wax layer on the top part of the leaf was removed by finger-wiping (right). Bar = 1 cm.

A total of 91.37 Gb clean reads (~166×) were produced using the Oxford Nanopore Technologies (ONT), complemented with a total of 35.17 Gb (~63×) of Illumina sequencing data. After genome assembly, polishing, and removal of haplotypic contigs, the final assembly size of the genome was 562.96 Mb, close to the flow cytometry-based estimate (Fig. S1), and the resulting contig N50 was ~1.60 Mb (Table S1). Nearly all (97.59%) of the Illumina sequencing data mapped to the genome assembly and 90.0% mapped properly oriented (Table S2). Genome completeness was evaluated via CEGMA v2.5 and BUSCO v 3.0 to be 96.07% and 93.49% respectively^31,32^ (Table S3).

From among the 1,614 plant-specific single copy orthologs in BUSCO database, 1509 (93.49%) were identified in the assembly, of which 1400 (86.74%) were complete and single copy (Table S3). Thus, a high quality genome sequence of *V. darrowii* was assembled.

### Chromosome-level assembly of Hi–C

To provide a chromosome-level assembly we then applied high-throughput chromosome conformation capture (Hi-C) sequencing, generating 32.81 Gb (60×) of 117.96 million Hi-C paired-end reads. Duplicate removal, sorting, and quality assessment were performed with Hi-C-Pro^35^. There were 55.37 million unique paired alignments to the genome, and 46.15 million pairs were valid for Hi-C scaffolding (Table S4). After correction of chromosome order and direction, the final assembly of *V. darrowii* genome was 563.0 Mb. After the Hi-C scaffolding, altogether 96.48% (543.12 Mb) of the assembly mapped to 12 pseudomolecules with 505.56 Mb being properly oriented (Table S5). The contig N50 of the final assembly was 1.50 Mb and scaffold N50 was 41.99 Mb, respectively (Tables S6). The total gaps was limited to 52.93 Kb (Table S6). The Hi-C heat map shows that 12 pseudochromosomes were well connected along the diagonal line (Fig. 2A). A high-quality chromosome-scale genome was assembled.

**Fig. 2.**
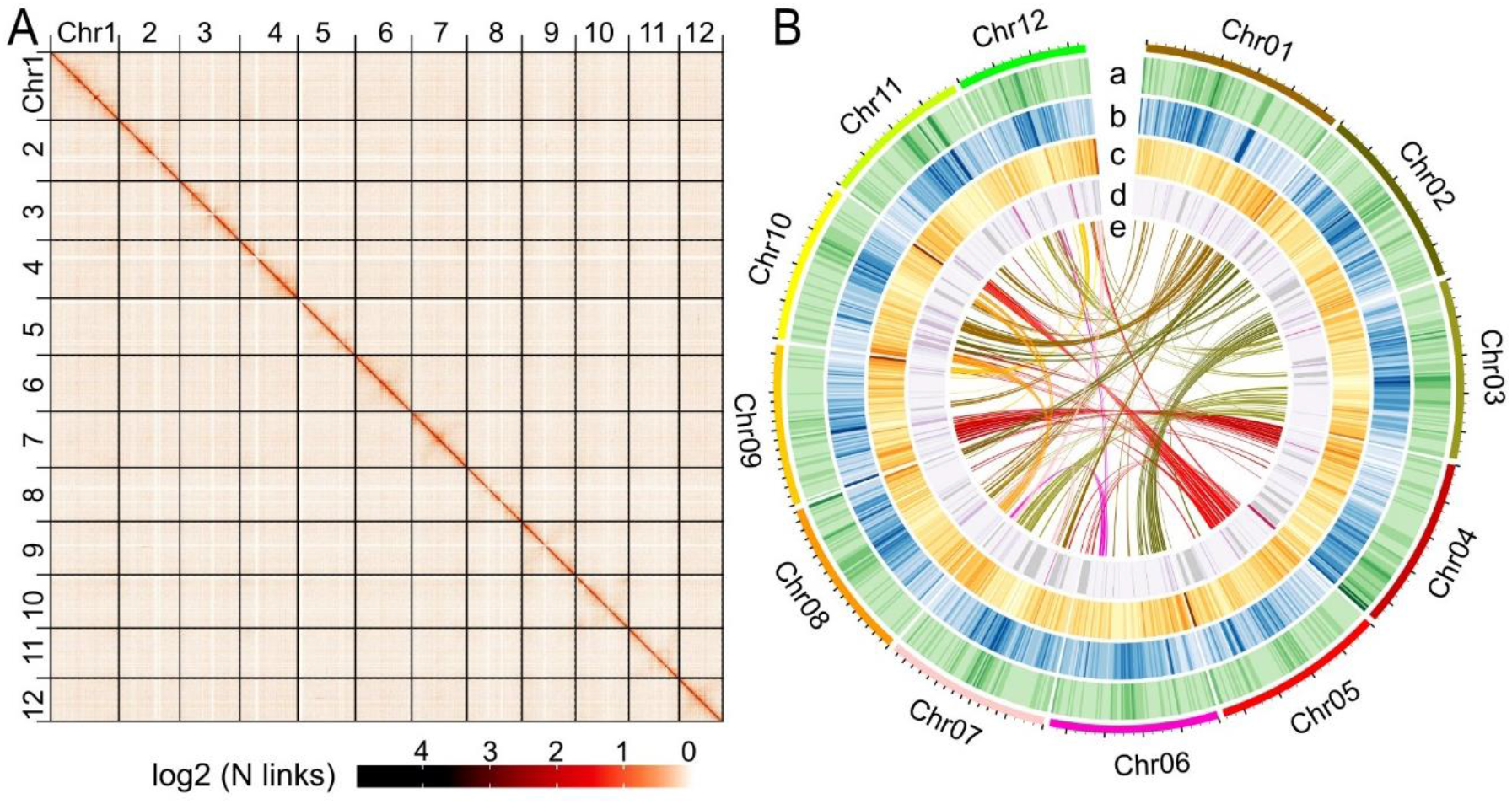
*V. darrowii* Genome assembly. (A) Hi–C heat map of chromosome interactions. (B) An overview of *V. darrowii* genomic features. Tracks: (a) GC content; (b) repeat sequence coverage; (c) gene density; (d) noncoding RNA density; (e) chromosome collinearity visualized by linking syntenic genes.

### Gene prediction and functional annotation

A combination of plant protein homology mapping, transcriptome data, and *ab initio* gene prediction was used to generate gene model predictions (Fig. S3, Table S7), and the different evidence tracks were combined to consensus gene models using EVidence Modeler^46^, leading to a total of 41 815 protein-coding genes (Table S8). The average gene length was 5076.09 bp, and the mean exon number of each gene was 5.02 (Table S8). In addition, 992 noncoding RNAs, including 119 microRNAs (miRNAs), 257 rRNAs and 616 tRNAs were detected (Table S8). The *V. darrowii* genome contained 333.76 Mb of repetitive sequence, accounting for 59.28% of the genome. Long terminal repeat (LTR) retrotransposons accounted for 24.47% of the genome, including 9.1% Copia-LTRs and 15.14% Gypsy-LTR (Table S9).

Through BLAT and GeneWise software^44,71^, 5300 pseudogenes were found (Table S8). Functional annotation was carried out using public databases, including GO (37.81% of genes were annotated), KEGG (26.45%), KOG (41.32%), Pfam (63.7), Swissprot (53.03), TrEMBL (79.92%), and Nr (80.18%) (Table 1). Only about 18.9% of the genome sequence was unannotated.

**Table 1.** Database and the corresponding annotated genes.

| Annotation database | Annotated number | Percentage (%) |
| --- | --- | --- |
| GO Annotation | 15,855 | 37.81 |
| KEGG Annotation | 11,092 | 26.45 |
| KOG Annotation | 17,326 | 41.32 |
| Pfam Annotation | 26,707 | 63.7 |
| Swissprot Annotation | 22,236 | 53.03 |
| TrEMBL Annotation | 33,511 | 79.92 |
| Nr Annotation | 33,619 | 80.18 |
| All Annotated | 34,005 | 81.1 |

### Comparative genomics analysis

To analyze gene family evolution and to obtain a phylogeny, we used Orthofinder and identified unique and shared gene families among blueberry and nine other species: *V. darrowii*, *V. corymbosum*, *V. macrocarpon*, *A. chinensis*, *A. thaliana*, *A. trichopoda*, *C. clementina*, *C. sinensis*, *F. vesca* and *S. lycopersicum*. All species shared 3732 gene families (Fig. S4), and altogether 347 gene families were specific to *V. darrowii* (Table S10). A phylogenetic tree was constructed using 1059 single-copy gene families, and calibrated using fossil data of *V. corymbosum* vs. *A. chinensis* (52-96 MYA), *V. macrocarpon* vs. *A. chinensis* (52-96 MYA), *V. corymbosum* vs. *A. trichopoda* (173-199 MYA), *C. clementina* vs. *A. thaliana* (96-104 MYA), *S. lycopersicum* vs. *C. sinensis* (99-114 MYA) as calibration points. The result indicated that *V. darrowii* and *V. corymbosum* diverged 4-25 MYA, preceded by split from *V. macrocarpon* at 7-39 MYA (Fig. 3A). Using CAFE^63^, we predicted altogether 232 contracted and 202 expanded gene families (Fig. 3B).

**Fig.3.**
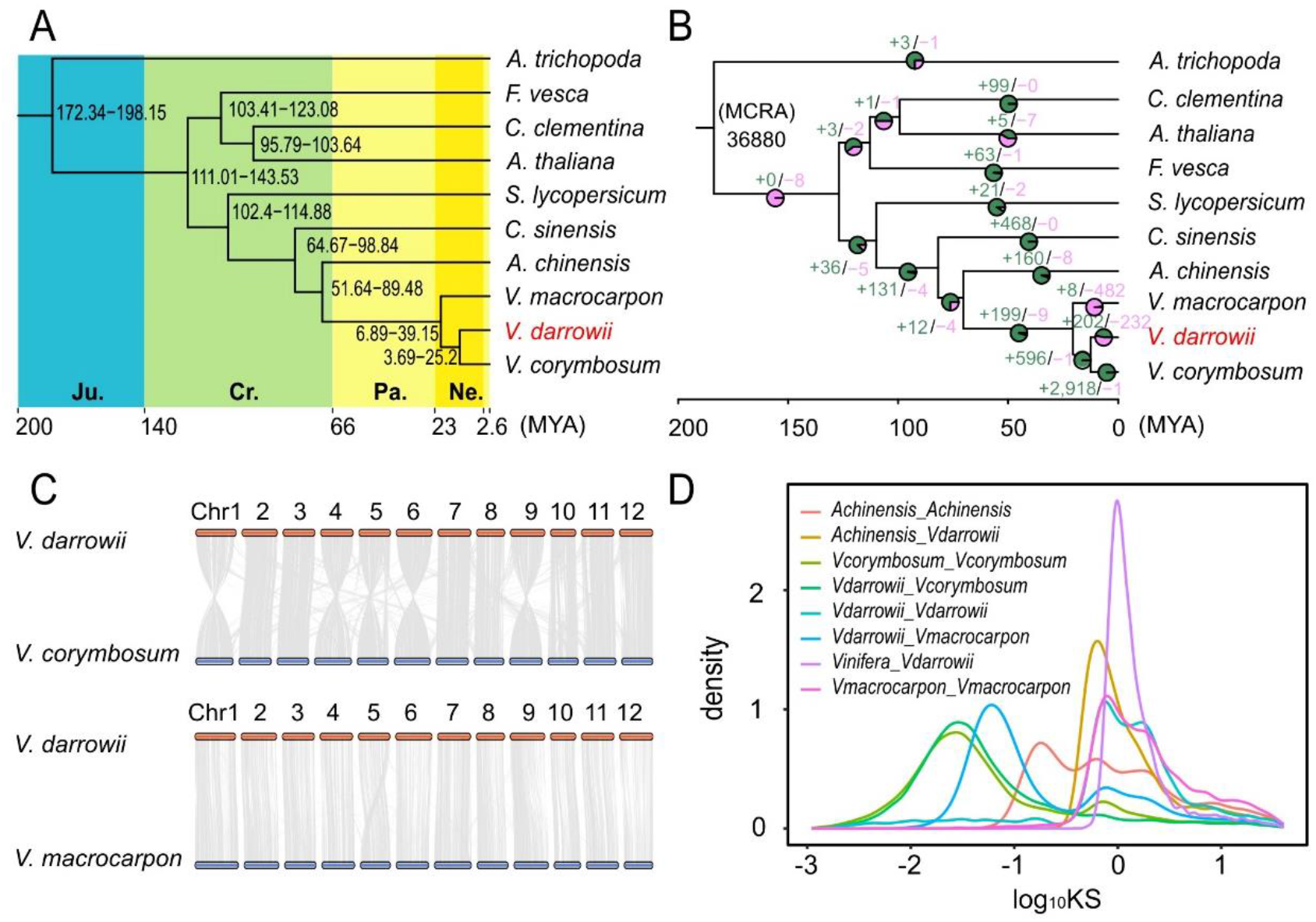
Genome comparison and evolution analysis. (A) Phylogenomic analysis with ten species and 1059 single-copy gene families. The divergence time was estimated with fossil times in TimeTree (http://www.timetree.org/).The divergent time was shown in million years after (MYA) (B) Gene family expansion and contraction in *V. darrowii*. The numbers of contracted gene family were in light green while the expanded gene family in light pink. (C) Synteny conservation between *V. darrowii* and two *Vaccinium* species *V. corymbousm* and *V. macrocarpon*. Altogether 96 334 genes from *V. corymbosum* and 28 527 genes from *V. macrocarpon* were collinear with *V*. *darrowii*. (D) Distribution of the synonymous substitution (Ks) rate for collinear genes from *V. darrowii*, *V. corymbosum*, *V. macrocarpon*, *A. chinensis* and *Vitis vinifera*.

### Collinearity and WGD analysis

Paralogous genes between *V. darrowii*, *V. macrocarpon* and *V. corymbosum* were identified via genome collinearity analysis. Genome organization between these species was well preserved, as chromosomes showed high levels of synteny (Fig 3C, Table S11, Table S12). Altogether 96 334 genes from *V. corymbosum* were collinear with *V. darrowii*, whereas 28 527 genes from *V. microcarpon* were syntenic. The analysis of synonymous substitutions (Ks) using syntelogs within each genome (Table S13), showed two whole genome duplication (WGD) events in *V. darrowii* (Fig. 3D). Syntenic comparison against grape (*Vitis vinifera*), a species that has undergone only the paleohexaploid (γ) event, showed one WGD event after divergence (Fig. S5), confirming that the older event is the shared γ event. The more recent WGD coincided with the Vm-α duplication reported earlier in cranberry^15^, and is a shared event since the species diverged much more recently (Fig. 3D). *V. corymbosum* on the other hand showed four-fold synteny, most likely resulting from a second WGD event when the tetraploid formed after divergence from *V.darrowii*. Thus, the three *Vaccinium* species share a common WGD event in addition to the ancient paleohexaploid WGT (γ) event approximately 130–150 Mya^72^, shared by all eudicots. Based on the divergence time of *V. darrowii* vs. *A. chinensis* (~78.5 MYA) and the Ks peak at 0.46, we estimated that the recent WGD event occurred at ~11.8 MYA (T = Ks/2r) in *V. darrowii*.

### Genome evolution in *V. darrowii*

As illustrated by the phylogeny, cranberry separated from blueberry first, followed by the split to *V. corymbosum* and *V. darrowii* (Fig. 3). Cranberry and *V. corymbosum* are both native to temperate regions, while only *V. darrowii* is a subtropical plant, suggesting temperate habitat to be the original environment of the common ancestor. In order to study the gene family evolution related to adaptation to subtropical climate, we analyzed gene family expansions and contractions between *V. darrowii* and *V. corymbosum*. KEGG pathways related to anthocyanin biosynthesis, such as ‘phenylpropanoid biosynthesis’, ‘flavonoid biosynthesis’, and ‘anthocyanin biosynthesis’ were contracted in *V. darrowii* (Fig. 4A). Anthocyanins play a protective role in plants during cold temperatures^73^, whereas high temperatures repress anthocyanin production in plants^74–76^. Thus, the contraction of anthocyanin-related gene families might be an adaptation result of *V. darrowii* to the warmer temperatures in the subtropics. From the individual gene families, the number of genes coding for Sn-2 acyl-lipid omega-3 desaturase was reduced from eight in *V. corymbosum* to one in *V. darrowii* (Supplemental Table 14). Sn-2 acyl-lipid omega-3 desaturase is a positive regulator of plant tolerance to cold temperatures and its knock-out mutant in *Arabidopsis* exhibits a chilling sensitive phenotype^77,78^. Unsaturated lipids can stay in liquid phase at lower temperatures than the saturated, allowing plants to maintain physiological functions during chilling stress^77^. Thus, the contraction of lipid desaturase in *V. darrowii* is a possible adaptation to the subtropical climate. Other terms enriched among the contracted gene families included ‘cell wall’, ‘root development’ and ‘iron ion binding’ (Fig. 4B). This might be related to the smaller size of *V. darrowii* in comparison to other *Vaccinium* species^19^.

**Fig. 4.**
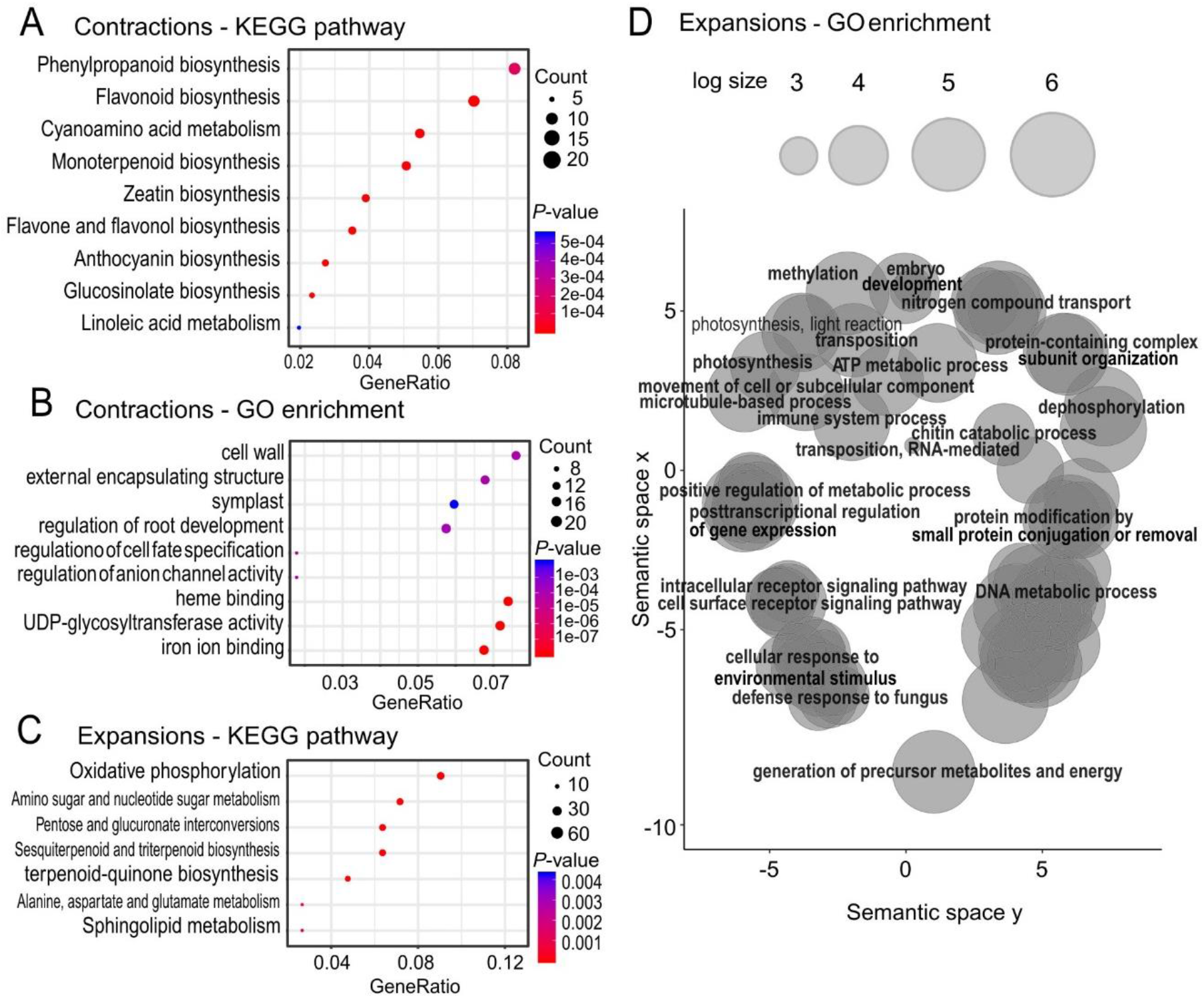
Functional enrichment analysis of expanded and contracted gene families. (A) KEGG pathway enrichment of the contracted gene families. (B) Gene ontology (GO) enrichment of the contracted gene families. (C) KEGG pathway enrichment of the expanded gene families. Gene Ratio represents the ratio of the number of the input genes annotated in a GO term to the number of all genes annotated in the term. The size of the dots represents the number of corresponding genes. (D) The expanded gene families were subject to GO enrichment analysis. A summarization of the genes in the enriched GO terms with Revigo was performed to see the relations between enriched GO terms.

Many of the GO terms enriched among the expanded gene families in *V. darrowii* were initially associated with chloroplasts and placed in contigs not mapped to the pseudochromosomes. After removal of the possible plastid contamination, secondary metabolism was enriched among KEGG annotation, including ‘amino sugar and nucleotide sugar metabolism’, ‘sesquiterpenoid and triterpenoid biosynthesis’ and ‘terpenoid-quinone biosynthesis’ (Fig. 4C). Plant secondary metabolism is one of the main adaptive mechanisms for coping with altered environmental conditions and biotic stressors^79,80^. To gain an overall view of the enriched GO terms among expanded gene families, we summarized the terms using Revigo^81^, and identified two larger clusters of terms with high similarity. Within the cluster defined by GO term ‘DNA metabolic process’, a sub-term ‘DNA repair’ was of the lowest *P*-value (Fig. 4D; Table S14). Genes associated with this term were mainly DNA helicases (Table S14). DNA repair systems and DNA helicases play an active role in plant thermotolerance^82^. Therefore, an expansion of the DNA helicases may contribute to adaptation to elevated temperatures. The ‘blue light signaling pathway’ was among the enriched GO terms associated with the cluster linked to environmental adaptation (Fig. 4D). The expanded gene family contained four duplicates of *Maintenance of Meristems* (*MAIN*) and *MAIN-like 1* genes (Table S14). MAIN and MAIN-like 1 interact with protein phosphatase PP7-like^83^ to regulate transposable element silencing, as well as chloroplast development and abiotic stress tolerance^84^. PP7L is closely related to PP7, a component in cryptochrome-mediated blue-light signaling^85^ and a regulator of the perception of red/far red light through the control of the phytochrome pathway^86^. The expansion of MAIN and MAIN-like 1 genes may therefore be due to altered light conditions in the subtropical regions and suppression of transposable element expression resulting from environmental stresses.

Interestingly, PsbP, a membrane extrinsic subunit of photosystem II ^87^, was among the 13 genes under positive selection in *V. darrowii* (Table S15). PsbP is essential for the stabilization of photosystem II, and decreased expression of *PsbP*, but not another extrinsic subunit gene *PsbQ*, resulted in pale-green-colored leaves and enhanced sensitivity to high light in tobacco^88^. Enhanced PsbP expression in *V. darrowii* may help to maintain photosynthesis in the drought/hot summer of the subtropical region. In conclusion, the expanded and contracted gene families reflect the adaptation of *V. darrowii* to subtropical climates.

To look for short term evolutionary adaptations, we carried out tandem duplication analysis with both *V. darrowii* and *V. corymbosum* (Table S16 and S17). After *P*-value adjustment with Bonferroni correction, the tandemly duplicated genes of *V. darrowii* and *V. corymbosum* were enriched for 131 and 468 GO terms, respectively (Fig. 5A). Altogether 55 terms were *V. darrowii* specific (Fig. 5A). Consistent with tandem expansion analyses in other species the enrichments were related to secondary metabolism ‘drug transport’, defense responses ‘response to fungus’ or adaptation to the environment ‘regulation of response to stress’ (Fig. 5B; Table S18). Overall the tandem expansions suggested adaptation to a new environment and genome rewiring of defenses against new pathogens. Interestingly, five *ERdj3B* were tandemly duplicated in *V. darrowii* (Table S18). *ERdj3B* encode a dnaJ domain protein that binds to heat shock proteins^89^. ERdj3B is essential for pollen development at high temperatures, and the corresponding *Arabidopsis* mutant produced few seeds at slightly elevated temperature, from 22 to 29 °C^89^. Fertilization in *Vaccinium* species is sensitive to high temperature^90,91^. The duplicated *ERdj3B* genes could indicate an evolutionary strategy for successful fertilization of *V. darrowii* in warmer regions.

**Fig. 5.**
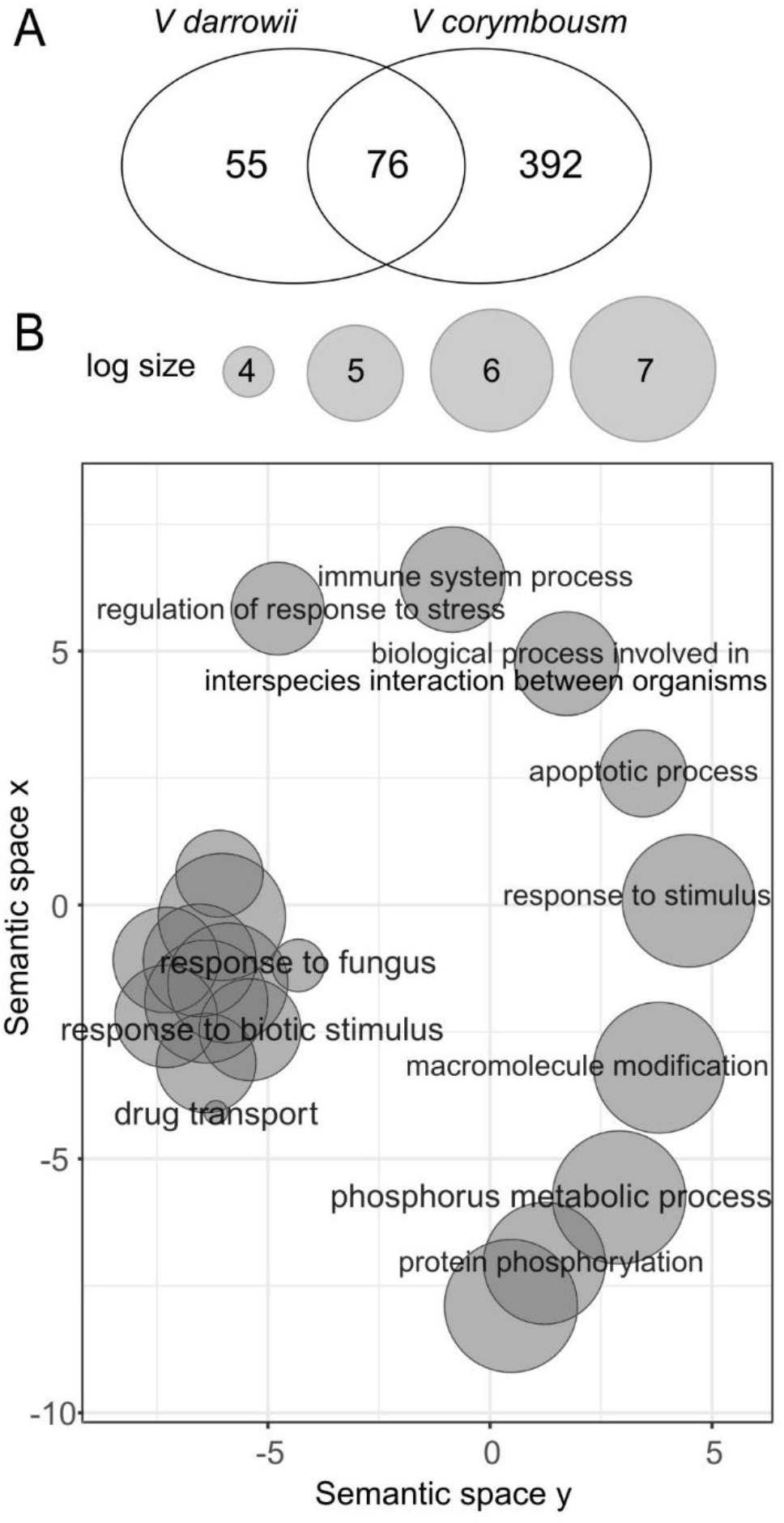
GO enrichment analysis of tandem duplicated genes. (A) Venn diagram showed the numbers of GO terms enriched in the tandem genes of *V. darrowii* and *V. corymbosum*. (B) The *V. darrowii*-specific GO terms were summarized with Revigo to see their relations.

### Flowering related genes in *V. darrowii*

Except for the florigen gene *FT*, other flowering regulatory genes in blueberry were rarely identified at the genome level^92,93^. *V. darrowii* exhibits a short vegetative phase, and flowering can occur already within one year^17,18^. Four *FPA* genes and six *Suppressor of Overexpression of CO 1* (*SOC1*) were among tandem duplications from the set of 26 genes assigned to the GO ‘regulation of flower development’ (Table S18). In *Arabidopsis*, FPA promotes early flowering via regulation of RNA processing which leads to reduced expression of the flowering-suppressor *FLC*^94–97^. The function of FPA in perennial plants has not been reported. SOC1 is major player in promoting flowering in many annual plants and several perennial plants, such as bamboo^98,99^. SOC1 directly promotes the expression of genes initiating flower development^100,101^. In trees, *SOC1* lowers the chilling requirement and promotes flower bud release from dormancy^17,102,103^. One of the major contributions of *V. darrowii* to blueberry breeding is the reduced chilling exposure time required by northern highbush blueberry for flowering, which resulted in the generation of the southern highbush blueberry^10,18^. Thus, we focused on *SOC1* and its tandem duplications. To confirm the homology relationships, we carried out multiple sequence alignment of the orthogroups containing genes annotated as SOC1 homologs and estimated a gene tree from the alignment. Closer inspection revealed a split into three clades of AGAMOUS-like paralogs: FYF clade, AGL14/19 clade and SOC1 clade, each clade named after the *Arabidopsis* homolog within the clade (Fig. 6A). The most distant clade, FYF, controls flower senescence and abscission in *Arabidopsis*^104^. In the AGL14/19 clade, AGL14 is expressed in roots in *Arabidopsis*, and therefore not likely involved in cold response of flowering, whereas variation in AGL19 has been found to be associated with vernalization response and timing of flowering^105^. Therefore, blueberry homologs in this clade may also be regulated under cold stress and could be regulators of flowering.

**Fig. 6.**
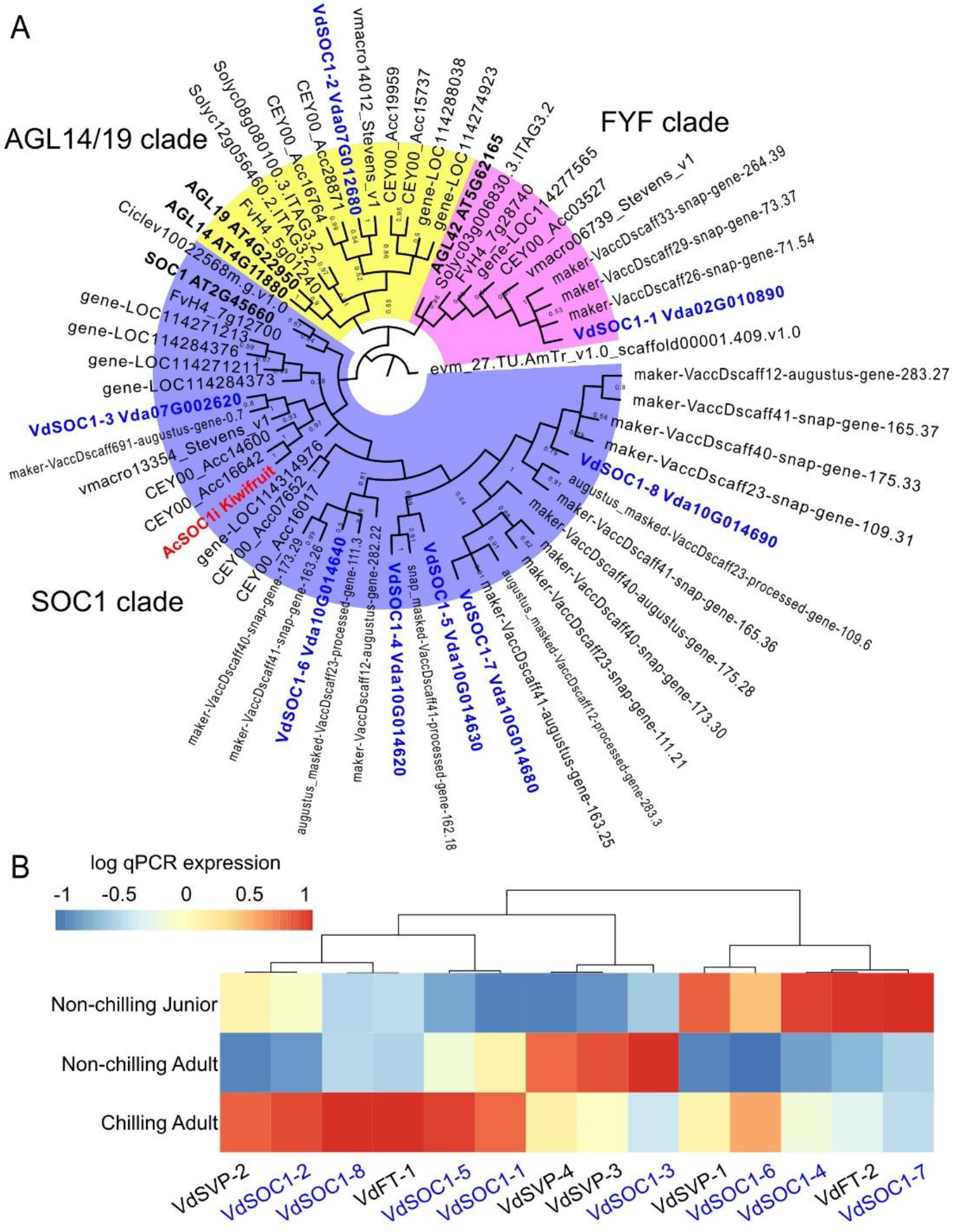
Phylogeny analysis and expression profile of the *SOC1* like genes in *V. darrowii*. (A) The SOC1 proteins were identified from an orthogroup that includes the *Arabidopsis SOC1* gene. Phylogenetic analysis of *SOC1*-like genes from *V. darrowii, V. macrocarpon* and *V. myrtillus* were performed using the complete protein sequences and clustered into three clades named with the corresponding *Arabidopisis* genes. The Maximum-Likelihood (ML) tree was reconstructed using MUSCLE alignment and MEGAX software. 500 bootstrap replicates were used to assess tree reliability. The tree was rooted with *Amborella trichopoda SOC1* gene. (B) Expression of *VdSOC1*s, and other flowering regulatory genes *VdFT*s and *VdSVP*s were examined in adult and junior clones of *V. darrowii*. A chilling treatment under natural conditions was conducted to induce flowering in the adult clones. The age and chilling effects on the expression were examined via qPCR using *VdTub.alpha3, VdTub.beta8* and *VdActin3* as references. The *VdSOC1*s were highlighted in blue characters.

Within the SOC1 clade, two typical SOC1, AtSOC1 from *Arabidopsis* and AcSOC1i from kiwifruit exhibited a split corresponding to plant phylogenetic relationships (Fig. 6A). AcSOC1i is an experimentally verified SOC1 gene in kiwifruit, promoting dormancy release^106^. The gene tree placed Vda07G002620 in the same clade together AcSOC1i, suggesting a possible similar function of these genes (Fig. 6A). The tandem duplicates including Vda10G014620, Vda10G014630, Vda10G014640, Vda10G014680, and Vda10G014690 grouped together, indicating their recent diversification (Fig. 6A).

To identify the orthology relationships, we investigated the expression pattern of all putative *SOC1* homologs together with other two key flowering-regulatory genes, the flowering promoting genes *Flowering Locus T* (*FT*) and the flowering suppression gene *Short Vegetative Phase* (*SVP*)^107,108^. We studied a total of eight *SOC1*s, two *FT*s and four *SVP*s annotated in *V. darrowii* genome (Table S19). With qPCR we examined leaf samples from two-year adult plants with or without chilling treatment, and also four-month old junior plants. We used the initiation of flower buds as a standard for a successful chilling treatment. The transcript level of *VdFT1* was high only in the chilling-treated plants while *VdFT2* was high in the junior plantlets (Figure. 6B). Thus, *VdFT1* could be the florigen of *V. darrowii*. *VdSOC1* −1 (Vda02G010890), −2 (Vda07G012680), −5 (Vda10G014630) and −8 (Vda10G014690) exhibited similar expression patterns to *VdFT1*, with highest transcript levels in plants with chilling treatment (Figure 6B). Thus, these four *VdSOC1* may function as the genes promoting flower development. The transcript levels of *VdSVP* −3 and −4 were highest in the non-chilling treated adult plants, and in the chilling-treated adult plants the transcript levels were much lower, indicating that *VdSVP* −3 and −4 were more likely the functional *SVP*s in *V. darrowii* (Figure 6B). Overall, we identified the key flowering regulatory genes in *V. darrowii*.

To test whether *SOC1* are tandemly duplicated among *Vaccinium* species, we analyzed genomes of two species from the other section of *Vaccinium*, cranberry and bilberry. There were five *SOC1* in *V. macrocarpon* with three being tandemly duplicated and seven in *V. myrtillus* with four being in tandem duplications (Fig. 6A). The numbers of both duplicated and total SOC1 gene models were comparable to *V. darrowii*. Thus, *SOC1* duplication is a universal in *Vaccinium*. This indicated that SOC1 genes may function in an unstudied complex pattern across the genus.

### Gene expression patterns between leaf, berry flesh and berry skin

The ‘powdery blue’ color is one of the major quality-determinant factors of blueberries in the market. The color depends on the content of anthocyanin pigments and wax, both of which are synthesized in the epidermal layer of the berry skin^109,110^. Interestingly, the leaf surfaces of *V. darrowii* are covered also by a thick layer of waxy cuticle, resembling the berry surface (Fig. 1). We performed RNA-seq analysis with samples from three different tissues, berry skin, flesh and leaves. Three biological repeats showed distinct clustering in the principal component analysis (Supplementary Fig. S5). Genes with tissue-specific expression were collected and visualized via a heatmap (Fig. 7A). There were 4131 genes specifically expressed in the leaf, and according to the Gene-ontology (GO) enrichment analysis they were mostly enriched for photosynthesis associated processes (Fig. 7B and C). This suggested accurate analysis, as photosynthesis is usually removed in the ripening blueberries. In the skin, 661 genes were specifically expressed (Fig. 7B), with GO enrichments for both cuticle and anthocyanin associated terms (Fig. 7C). The key steps of cuticle biosynthesis were included in the enriched GO terms: ‘citrate lyase complex’ and ‘ATP citrate synthase activity’. These are involved in the conversion of citrate to acetyl-CoA; acetyl-CoA participates in ‘fatty acid derivative biosynthetic process’ via ‘transferase activity, transferring acyl groups’; then fatty acids are subjected to ‘fatty acid elongation’ in a series of ‘fatty acid elongase complex’, and then ‘cutin/wax biosynthesis process’ is initiated (Fig 7C). Anthocyanins are synthesized from phenylalanine ammonia with trans-cinnamate and flavonoids as intermediates^111,112^. Accordingly, the GO terms ‘phenylalanine ammonia−lyase activity’, ‘trans−cinnamate 4−monooxygenase activity’ and ‘flavonoid biosynthetic process’ were significantly enriched among the skin-specific genes (Fig. 7C). Altogether, the strategy of using skins to identify cuticle related genes was successful. Among 425 flesh-specific genes, common processes involved in berry ripening ^113^, including ‘plant-type cell wall loosening/modification’ and ‘hormone metabolic process’ (Fig 7B and C), were enriched.

**Fig.7.**
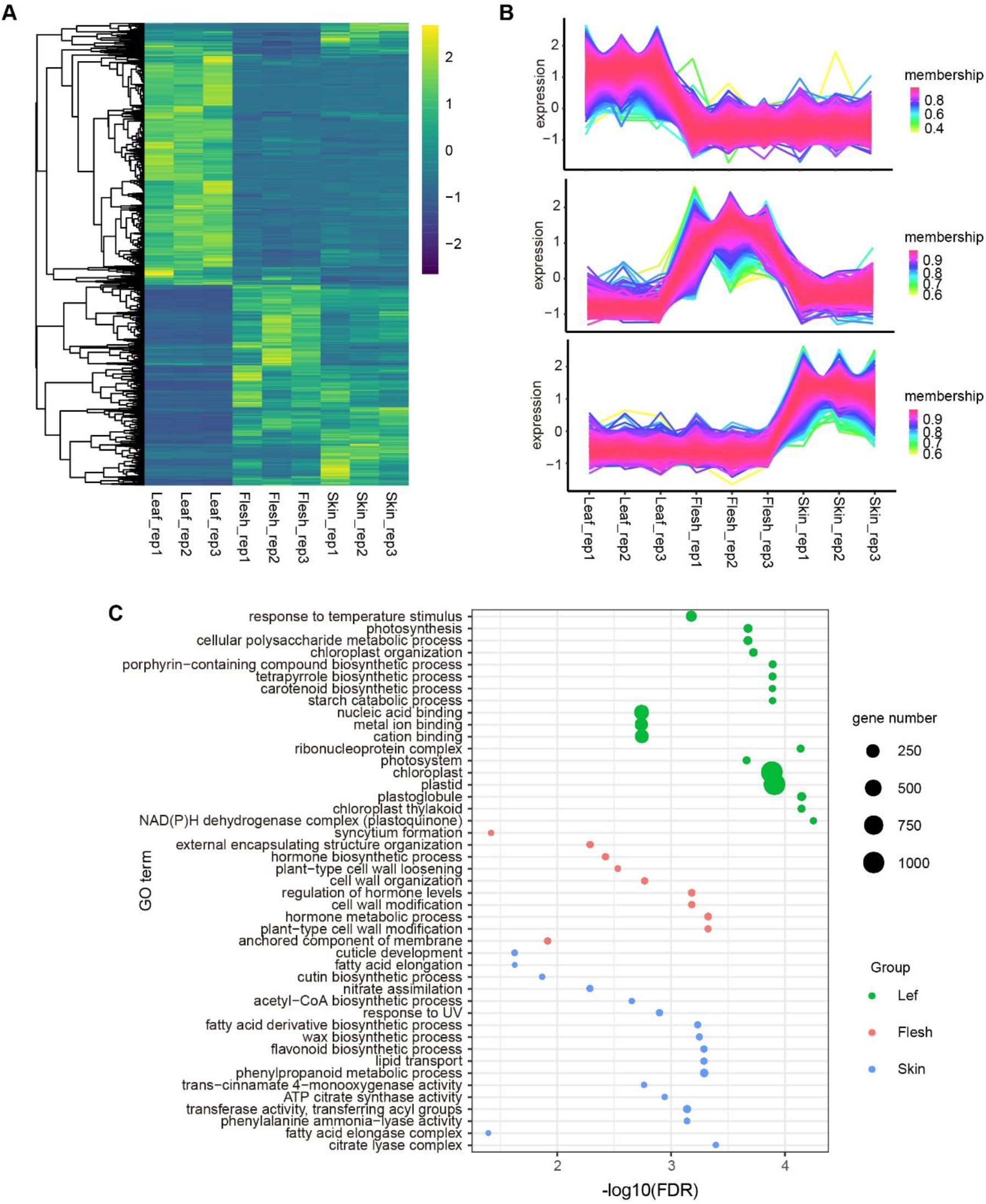
Tissue specific gene profiles in *V. darrowii.* (A) Heatmap of DEGs expression between three tissue leaf, flesh and skin. (B) Gene expression line graph of genes specifically expressed in leaf, flesh or skin. Membership values in color indicates the degree of the genes belonging to the cluster(C) GO enrichment of the three group tissue specific genes. The size of the dots represents the number of corresponding genes.

### Cuticle biosynthesis pathway

Among the two important functions of blueberry skins, the anthocyanin synthesis pathway has been well documented^14,114^. However, the wax biosynthesis pathway is not well understood, although several berry genomes have been published. Cuticle, consisting of wax and cutin, not only directly influences the color of blueberries, but also has a direct influence on the storage loss^109,115–117^. We identified the cuticle synthesis genes in *V. darrowii* from the Orthofinder clustering to their known *Arabidopsis* orthologs (Table S20). Interestingly, many genes related to wax synthesis showed small-scale tandem duplications (Table S17 and 20). Notably, 4-7 duplicates were found for *Eceriferum 1* (*CER1*) and *CER3*, two key synthesis enzymes of the wax monomer alkane ^118,119^, and *Mid-chain Alkane Hydroxylase 1* (*MAH1*) and *Wax Ester Synthase/Acyl-CoA:Diacylglycerol Acyltransferase* (*WSD1*), which function in the last two steps of wax biosynthesis in plants^120,121^ (Table S17 and 20). These duplications may explain the enhanced wax accumulation on blueberries.

We next explored the complete pathway of cuticle biosynthesis using the predicted relevant homologs in *Arabidopsis* (Table S20) and plotted them together with the expression levels between samples of leaf, flesh and berry skins (Fig. 8). Cuticle biosynthesis in model plants is considered to be epidermis-specific^110^. Consistently, most genes listed in the pathway exhibited a higher expression in the epidermis samples including leaves and berry skins (Fig. 7). Cutin Deficient (CD) is the key cutin synthase at the cell wall for the final step of cutin formation in many land plants^122^. None of the homologs of *CD* gene in *V. darrowii* were expressed. The possibility of *CD* gene missed from the annotation was excluded by blasting the CD genes to the genome sequence of *V. darrowii*, and no other homologs were found. Thus, cutin synthesis in the cell wall might differ between *V. darrowii* and other plants studied earlier^122^.

**Fig. 8.**
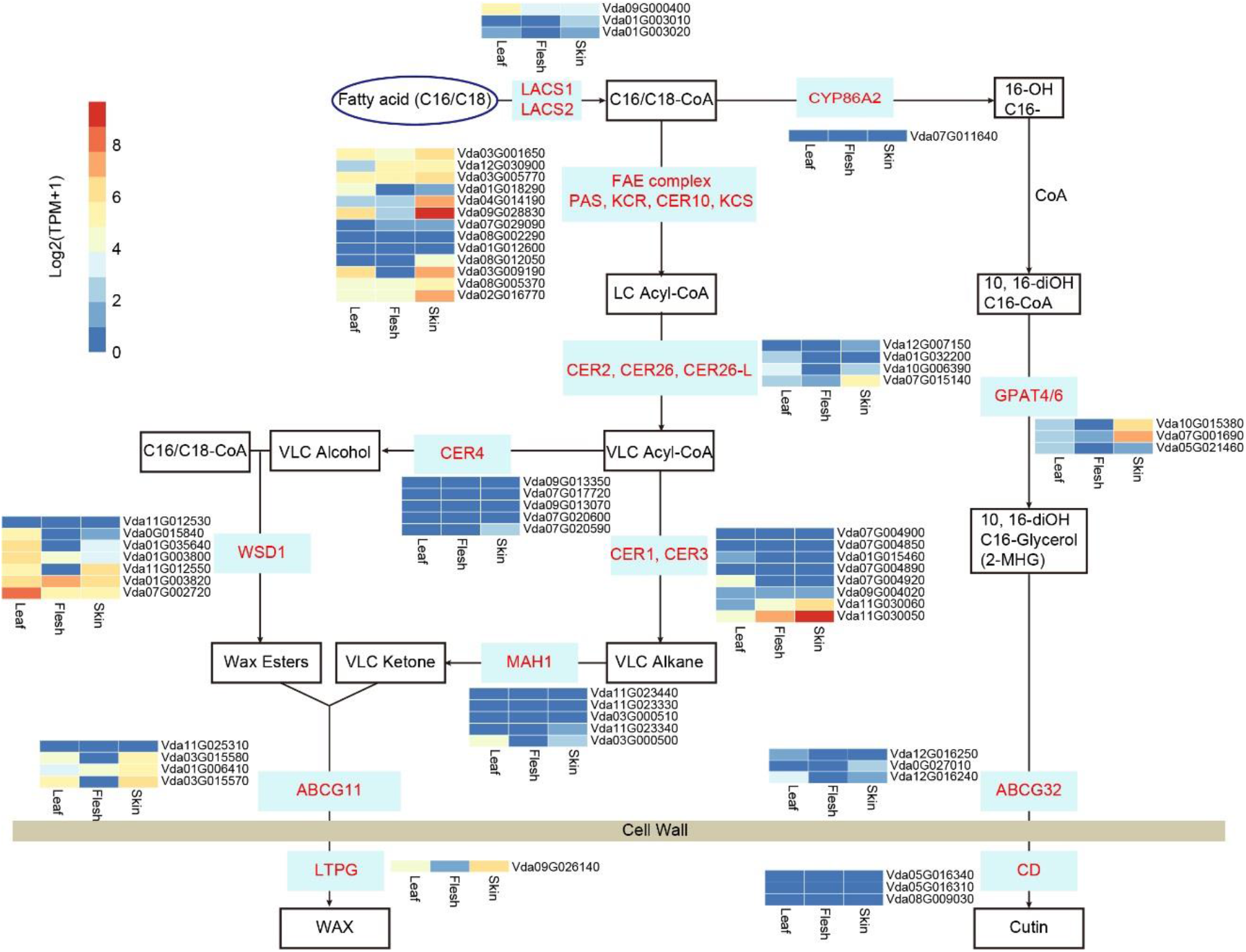
Putative cuticle pathway predicted in *V. darrowii* with transcriptome expression data. The genes in this pathway were identified using BLASTP alignment with *Arabidopsis* corresponding amino acid sequences. Transcript levels from RNAseq data were visualized in heatmap and normalized with log2(TPM+1).

## Discussion

*Vaccinium* species are widely distributed, from the tropical regions to the polar circle and tundra. *V. darrowii* is a wild blueberry species of practical and economical value. Since it was used to breed the subtropical adapted blueberry cultivars, studies on its unique traits including lowered chilling requirement, early flowering and heat tolerance are increasing^10^. The phylogenetic tree suggests that the *V. darrowii* originated in the temperate region and later on adapted to subtropical regions (Fig. 2 A). We identified several genome characteristics that possibly contributed to the adaptation of *V. darrowii* to subtropical climates.

Tandem duplicates contribute to gene family expansions and allow alternative transcriptional regulation, and thus have been considered a common strategy for adaptive evolution to altered environments^123,124^. A clear functional bias in the tandemly duplicated genes has been observed in plants, making tandem genes good index for environmental adaptation^123,125,126^. The GO enrichment analysis with the tandemly duplicated genes of *V. darrowii* provided an overview of possible adaptation traits. Defense related GO terms were specifically abundant in *V. darrowii* (Fig.5; Table S18). The same terms were also enriched in the gene family expansion analysis, indicating that the retention of these genes may assist the growth of *V.darrowii* in subtropical environments. Additionally, the duplications consisted of multiple genes that have been reported to be involved in regulation of plant tolerance to elevated temperatures, such as genes involved in secondary metabolism and response to stresses (Table S17 and 18). Notably, the genes mediating plant fertilization in warmer temperature were also present in the tandem duplicated genes (Table S17 and 18). It is common in farms that high temperature reduces fertilization of blueberry cultivars^91^. The tandem genes related to fertilization may help *V. darrowii* propagating in subtropical regions. In addition, 26 genes were predicted within the GO term ‘regulation of flower development’ in *V. darrowii* (Table S17 and 18). Since *V. darrowii* originated from high latitudes (Fig. 1), successful flowering and fertilization in the conditions with reduced chilling period and relatively hot temperature may be essential for the propagation *V. darrowii* at low latitudes.

SOC1 plays a key role in promotion of plant transition from vegetative phase to reproductive phase in annual plants^98,101^. In perennial species, SOC1 is a major regulator of bud dormancy release^17^. A role for SOC1 in regulating the requirement for an extended cold treatment to release flower bud dormancy has been reported in kiwifruit, Japanese apricot and sweet cheery^106,127,128^. The *SOC1* genes in *V. darrowii* were highly duplicated, and six of eight *VdSOC1*s were tandem duplications (Table S17). Although the eight VdSOC1 were split into three clades, six of them were clustered together with the *Arabidopsis* SOC1 and functional SOC1 in kiwifruit (Fig. 6A). This suggested that the VdSOC1s may function in flowering promoting and dormancy breaking activities. However, timing and location of gene expression largely determinate the function of genes. The Vda07G002620 (VdSOC1-3) shared the same sub-clade with the kiwifruit AcSOC1i, but chilling treatment suppressed its expression, suggesting it might be not the functional SOC1 in *V. darrowii* (Fig. 6). However, Vda07G012680 (VdSOC1-2) in another clade with the *Arabidopsis* AGL42, was expressed at the highest level at chilling treated plants (Fig. 6). It may function as a flowering promoter as the AGL42 was reported to promote flowering in *Arabidopsis*^129^. SOC1 was also reported to suppress flowering in strawberry^130^. Thus, complex or alternative roles might be played by *SOC1* genes with high sequence identities. In *Arabidopsis*, the transcript levels for *SOC1* in leaves firstly increased and then decreased during development^100^. This is mainly due to the complex function of *SOC1* in both vegetative phase and reproductive phase^131–133^. The single *SOC1* in *Arabidopsis* integrates complex signaling associated with pathways of thermosensor, autonomous, gibberellin, verbalization and photoperiod^134,135^. In *V. darrowii*, four tandem duplicated VdSOC1s in the same cluster exhibited different expression patterns (Fig. 6B), implying a functional differentiation between these highly similar genes. This hypothesis needs to be examined *in vivo* and is beyond the scope of this paper. Future studies could reveal the individual function of each *VdSOC1*. Our genome information here would facilitate the mining of these functional genes in the future.

Waxy surface is a very typical characteristic of *Vaccinium* berries. Multiple health-promoting activities of wax components have been discovered, such as triterpenoids which play roles as anticancer, anti-inflammation and antimicrobial agents^136^. Wax monomers vary between *Vaccinium* species and have been well documented both morphologically and chemically^116,137,138^. However, genes associated with wax biosynthesis have not been well defined in the published genomes of *Vaccinium*^14,15^. Here, we identified all candidate cuticle biosynthesis genes in *V. darrowii* (Fig. 8), which facilitate breeding of desired powdery color of blueberries. Compared to cutin, wax synthesis genes of *V. darrowii* exhibited multiple tandem duplicates (Table S17; Fig. 8). The tandem duplicates of wax synthesis genes was also observed in the northern highbush blueberry^14^. This suggested that enhanced wax synthesis commonly exists in all blueberries, which is consistent with the universal waxy berries from all blueberry cultivars^116,137,138^. One unique character of *V. darrowii* is its waxy leaves, which phenotypically resemble to the powdery surface of blueberries (Fig. 1B and C). Some cuticle synthesis genes were highly expressed in the leaf in comparison to berry skin (Fig. 8). Thereby, using leaves to study wax biosynthesis pathway instead of berries is possible in *V. darrowii*. This would dramatically shorten the time for functional examination of wax associated genes, as leaves are available much earlier than berries. Thus, *V. darrowii* may serve as a model plant of the *Vaccinium* genera for wax-related studies.

Blueberry is a relatively young crop species with very little breeding, in part due to the lack of genome-level information on the species. Similar to many other commercially important crop species, blueberry cultivars are of polyploid origin, being mostly tetraploids or hexaploids. Polyploid genomes may confer blueberries with heterosis-like advantages, yielding improved growth, stress resistance and berry size. However, the resulting complex genome structure make the polyploid species less amenable to basic research in comparison to diploid genomes. One way to gain molecular knowledge of the polyploid cultivars is to study the diploid progenitors. For example, in strawberry studies, the diploid woodland strawberry (*Fragaria vesca*) has been widely used as a model because of its simple genome and easy genome-editing characteristics^139–141^. However, no diploid blueberry species has been proposed as a model yet. *V. darrowii* is naturally diploid. With the genome sequence, it could emerge as a blueberry model for the subtropical species. Other advantages of *V. darrowii* as a model blueberry also include early flowering, self-compatibility, and smaller plant size. Recently, birch has been promoted as a promising model tree with a small genome size, and accelerated flowering within one year via high CO2 treatment^142,143^, and on the other hand, the ‘Mini-Citrus’, a model promoted for orange species, can bear 1-2 fruits in the first year^144^. These two woody perennial species show significant advantages compared to the most used model tree, poplar, which is dioecious and has very long vegetative phase^145,146^. We propose that *V. darrowii* could be not only a model plant for blueberry species, but also a model for subtropical woody species.

## Materials and Methods

### Growth conditions and plant material

*Vaccinium darrowii* Camp was obtained from Stephen F. Austin State University, Texas, USA by Prof. Hong Yu, Nanjing botanical garden Mem Sun Yat-sen, and was cultivated with a mixture of peat, vermiculite and bark pieces (1:1:1). Growth room conditions were 150-200 μmol m^−2^ s^−2^ light, 60% humidity, 12/12 h (light/dark) photoperiod and 23/18 °C (day/night). The outdoor growing area is in Hangzhou, a subtropical city in China. The seedlings from cuttings of a single clone of *V. darrowii* were used for this study.

### Flow cytometry examination

Nuclei of *V. darrowii* were extracted from young leaf tissues as previously described^20^ with modifications according to^21^. In brief, leaves submerged in ice-cold nuclei isolation buffer were chopped with a sharp razor, then the nuclei were collected and stained with DNA fluorochrome (propidium iodine, 50 mg/ml). Flow cytometric examination was conducted on the BD FACSCalibur flow cytometer (Becton Dickinson, San Jose, CA) with tomato (*Solanum lycopersicum*, genome size 950 Mb) as reference.

### DNA preparation and sequencing

Young leaves from a 3-year old single cutting of *V. darrowii* were collected for DNA extraction with CTAB method^22^. DNA was further purified via VAHTS DNA Clean Beads (N411-03, Vazyme, China) and then used for DNA sequence preparation ^23^.

For Illumina PE150 sequencing, purified DNA was fragmented via ultra-sonication for library construction (350 bp). A total 35.17 Gb clear reads (around 60×) were obtained with Q20 > 97%, Q30 > 92%. The possibility of DNA contamination was excluded via alignment examination. BLAST alignment of randomly selected 10, 000 reads returned as best matches only to the closely related plant species, *V. macrocarpon* (51.37%) and *Vitis vinifera* (3.82%)^24^. The proportion of plastid sequence was less than 0.02% which was confirmed by comparison with the blueberry chloroplast genome (KJ773964.1; 1 320 bp). The genome characteristics were assessed by K-mer analysis ^25^. The estimated genome size is 555.38 Mbp with 42.92 repeat sequence and 1.27% heterozygosity.

For Nanopore sequencing, gDNA was repaired with NEB Next FFPE DNA Repair Mix kit (M6630, USA), and the library was constructed with ONT Template prep kit (SQK-LSK109, Nanopore, UK). Large fragments were collected with the BluePippin system (Sage Science, USA) and pipetted into a R9 flow cell. Sequencing was carried out with Nanopore sequencing technology at Biomarker Co., Ltd. using ONT PromethION platform with Corresponding R9 cell and ONT sequencing reagents kit (EXP-FLP001.PRO.6, UK), a total of 98.1 Gb high quality reads, corresponding to a depth of 166×, were obtained^26^.

### RNA preparation and sequencing

The mature berries were separated into flesh and skin via manual pealing. RNA from flesh, skin, and fully expanded young leaves were used for tissue-specific expression assay, while RNA from branches, leaves, flowers and berries of the sequenced *V. darrowii* individual were used for gene annotation. RNA extraction was conducted according to^27^. Briefly, the isolated RNA (5 μg) was purified with VAHTS mRNA Capture Beads (N401-01, Vazyme, China) for two rounds, and then subjected to fragmentation treatment. The cDNA libraries were constructed with NEBNext^®^ Ultra™ RNA Library Prep Kit (E7530L, NEB, USA), and then sequenced with paired-end sequencing on an IlluminaHiseq4000 (LC Sceiences,USA) platform at LC-Biotechnologies (Hangzhou) Co., Ltd.

### Genome assembly

Long read error correction was carried out with Canu (https://github.com/marbl/canu, v1.5)^28^ using criteria ‘genomeSize=1000000000’ and ‘corOutCoverage=50’. Next, overlapping was carried out with a highly sensitive overlapper MHAP (mhap-2.1.2, option ‘corMhapSensitivity=low/normal/high’), and subsequent error correction was performed though Falcon_sense method. Additionally, WTDBG2 and Smartdenovo assemblies were carried out followed by three rounds of error correction using Racon and adjustment via Pilon^29,30^. The assembly quality was evaluated via alignment of the reads from short read sequencing, altogether 97.59% of the reads were mapped, with 90% mapping uniquely. Genome completeness was evaluated via CEGMA v2.5 and BUSCO v 3.0 to be 96.07% and 93.49%, respectively ^31,32^.

### Hi-C Assembly

Hi-C fragment libraries were constructed in sizes from 300-700 bp and sequenced with Illumina platform^33^. Raw reads were then subjected to removal of adapters and removal of low quality reads, and trimming of low quality bases. The trimmed reads, altogether 60x coverage of the draft genome, were truncated at the putative Hi-C junctions and then aligned to the assembly with BWA aligner^34^. The uniquely aligned reads with quality ≥ Q20 were kept for further analysis. Invalid read pairs were filtered out via HiC-Prov2.8.1 ^35^. The uniquely mapping read pairs totaled 46.94% of the clean reads, of which 83.35% were valid interaction pairs. The valid interaction pairs were clustered, ordered and oriented onto pseudochromosomes via LACHESIS ^36^. Twelve pseudo-chromosomes were constructed manually with 96.48% of valid interaction pairs anchored, and 93.08% were properly ordered and oriented.

### Repeat sequences, non-coding RNA and pseudogene analysis

For repeat sequences, LTR FINDER (v 1.05) and Repeat Scout (v1.0.5) were used to build a primary database of repeat sequences of *V. darrowii* with default parameters^37,38^. The database was classified with PASTEClassifier and then combined with Repbase databases^39,40^. RepeatMasker v. 4.0.5 was used to screen repeat sequences that includes simple repeats, satellites, and low-complexity repeats with parameter -nolow -no_is -norna -engine wublast -qq -frag 20000 ^41^. For non-coding RNAs, rRNA and microRNA were predicted with Rfam database; tRNA was predicted with tRNAscan-SE v1.3.1 with option “-E -H”^42,43^. Pseudogene homolog sequences were aligned via GenBlast A v1.0.4 and then the frame shifts and internal stop code were screened via GeneWise v 2.4.1^44,45^.

### Protein-coding gene prediction and functional annotation

Gene prediction was performed with EVM v1.1.1 pipeline^46^. Briefly, *de novo* gene prediction was conducted with Genscan^47^, Augustus v2.4 ^48^, Glimmer HMM v3.0.4 ^49^, Gene ID v1.4 ^50^ and SNAP v 2006-07-28 ^51^ with default parameters. GeMoMa v1.3.1 was used for homolog prediction ^52^. The RNA reads were assembled via Hisat v2.0.4 ^53^ and Stringtie v1.2.3 ^54^, and then TransDecoder v2.0 and GeneMarkE-T v5.1 were subsequently applied for gene prediction^55^. RNA transcript assembly and unigene sequence prediction were conducted via PASA v2.0.2 ^56^. The prediction methods above were combined in EVM 1,1.1 and revised via PASA v2.0.2. The annotation of protein-coding genes was performed via BLAST v2.2.31 ^24^ (-evalue 1e-5) against the NR^57^, KOG^58^, GO^59^, KEGG^60^ and TrEMBL^61^ databases.

### Genome comparison and evolution assay

The protein data of representative plant species, including *V. darrowii*, *V. corymbosum*, *V. macrocarpon*, *A. chinensis*, *A. thaliana*, *A. trichopoda*, *C. clementina*, *C. sinensis*, *F. vesca* and *S. lycopersicum* were retrieved for gene family clustering and phylogenetic analysis. The protein sequences were clustered into orthogroups by using the OrthoFinder software (version 2.4) with default parameter settings. The pairwise similarities between the proteins from all species were searched in an all-vs.-all manner with Diamond with an E-value ≤ 0.001. The single-copy families among the analyzed genomes were subjected for multiple alignment by using MAFFT v7.205, and the alignments were trimmed using Gblocks v0.91b. The aligned gene family sequences were then concatenated and a phylogenetic tree was constructed using maximum likelihood (ML) method by IQ-TREE v1.6.11, and bootstrap was set to 1000. *A. trichopoda* was used as the root for the tree.

The divergence time was estimated using MCMCtree, incorporated in the PAML v4.9i software^62^. The fossil times used for calibrating the tree were obtained from TimeTree (http://www.timetree.org/). Gene family expansions and contractions were analyzed with CAFE (v4.2) with default settings^63^.

### Synteny and Ks analysis

All-vs-all BLASTP analyses of proteins were performed between and within the selected species, and the results were filtered using cutoff value of E<10E-3. Then, BLASTP results and genome annotation files of each species were used to identify syntenic blocks in MCScanX with default parameters^64^. The synonymous mutation rate (Ks) values of the homologs within collinear blocks were calculated using the wgd v1.1.0 software. The values of all gene pairs were plotted to identify putative whole-genome duplication (WGD) events. The duplication time was estimated using the formula t =Ks/2r, which represents the neutral substitution rate, and was used to estimate the divergence time between blueberry and other species. Tandem duplications were obtained as part of the MCScanX analysis.

### Transcriptome analysis

The cDNA libraries of leaf, flesh, and skin samples were constructed for RNA-seq, and three biological replicates were performed for each tissue. Trimmomatic v0.39 was used to remove low-quality and adapter sequences to obtain clean data (Bolger et al. 2014). HISAT2 v2.1.0 (Kim et al. 2015) was used to align the clean reads to the reference genome. Gene model prediction and calculation of gene expression was carried out with StringTie v1.3.4 (Pertea et al. 2016). DESeq2 (Love et al. 2014) was used for differential expression analysis between tissues. *C*-means (Wu and Gu 2017) was carried out for the set of screened differentially expressed genes (DEGs) to find groups of genes with specific expression patterns in tissues. For the detection of DEGs between tissues, fold change > 2 and false discovery rate (FDR) < 0.01 were used as cutoff values. To understand the biological functions of those gene clusters, GO (Gene ontology) enrichment analysis was performed by GOATOOLS^65^. Gene expression in the samples was plotted by R package pheatmap.

### Chilling treatments and qPCR procedures

Two-year-old cuttings from one clone of *V. darrowii* were moved outdoors from a growth room in middle of September, 2019. This allowed *V. darrowii* to grow naturally in the subtropical city Hangzhou. All the cuttings were moved into growth room (23°C) in the beginning of November, except the ones left for chilling treatment. Chilling treatment was carried out outdoors until the middle December. The average temperature outdoor in November was 10-18°C, December was 5-12°C. The cuttings were moved into a growth room and kept for three weeks for flowering induction. Cutting individuals that successfully developed flower buds were used for leaf collections. The non-chilling treated two-year-old and four-month-old cuttings of *V. darrowii* were collected at the same time. RNA was extracted using RNAprep Pure kit (DP441, Tiangen, China) for RNA isolation, and then the cDNA was synthesized with HiScript^®^ III 1st Strand cDNA Synthesis Kit (R312, Vazyme, China). Primers were designed with Real-time PCR (TaqMan) Primer and Probes Design Tool from GenScript wesite (www.genscript.com/tools/real-time-pcr-taqman-primer-design-tool) and listed in Supplemental Table 21. qPCR was applied as described in^66^ with Bio-Rad CFX touch. The raw cycle threshold values were processed in Qbase+ 2.1 ^67^ using *Tubulin α2*, *Tubulin α3* and *Tubulin β8* as reference genes.

### Data availability

Genome sequence and analysis associated data were deposit on CoGe (https://genomevolution.org/coge/GenomeInfo.pl?gid=62137). Clear reads from RNA-seq were uploaded to SRAs of NCBI with a deposit number PRJNA758745 (https://dataview.ncbi.nlm.nih.gov/object/PRJNA758745?reviewer=67gtdiagbfut5pmvf1vjigeu5v)

## Acknowledgements

These studies were supported by the National Natural Science Foundation of China (Grant No. 31700224); the Innovative Scientific and Technological Talents in Henan Province (No. 20HASTIT041) and Outstanding Youth Foundation of Henan Province (No. 202300410041), as well as Nanyang Technological University startup grant and Academy of Finland (decisions 318288 and 319947) to J.S. We gratefully acknowledge Prof. Wenwu Wu (Zhejiang A and F University) for the help on RNA-seq analysis, Dr. Xiaohong Zhou for all the help on plant cultivation. No conflict of interest declared.

## Author Contributions

FC designed the experiments. FC and JS conceived and interpreted the data. XY and JS performed the bioinformatic analysis. FC, XL performed the experiments. FC and ZH acquired funding. FC, XY and JS wrote the first manuscript, which all authors edited and approved. None conflict of interests were declared.

## Supplemental Figures

**Supplemental Figure 1.**
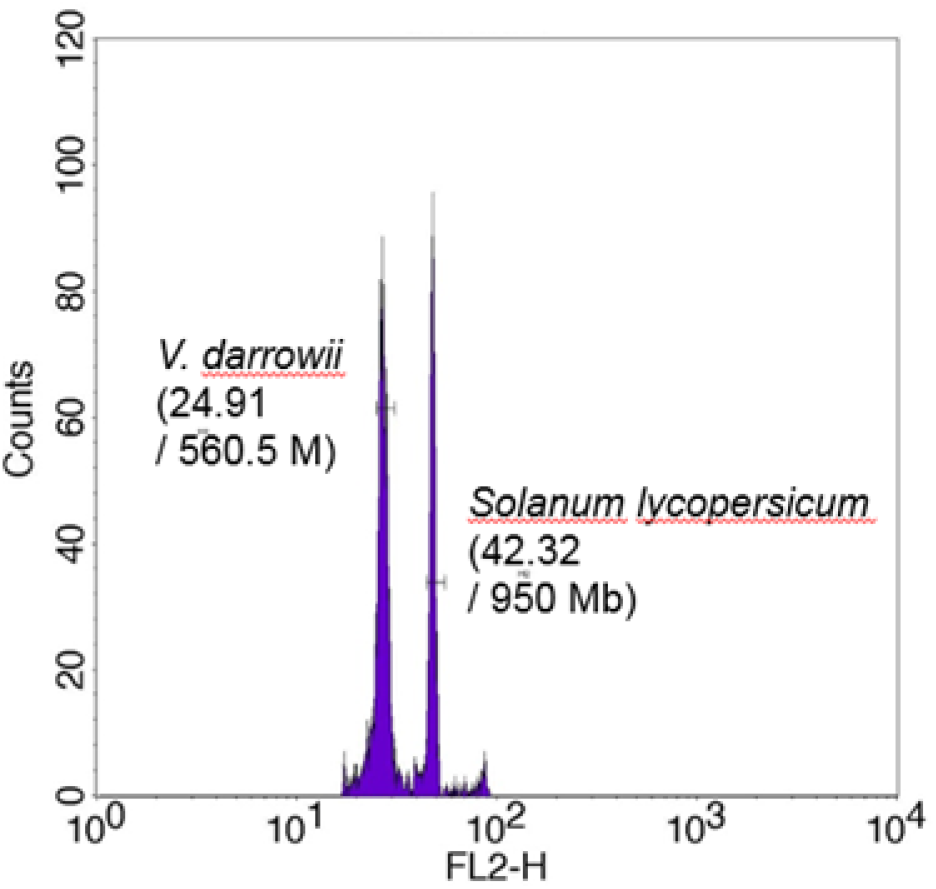
Flow cytometric analysis. The genome size of *Solanum lycopersicum* is 950 Mb, and the peak value is 42.32. The peak value of *V. darrowii* is 24.91, with ratio 0.59 in comparison with *Solanum lycopersicum*. The genome size of *V. darrowii* was estimated to be 560.5 Mb. The flow cytometry assay were performed three times with all the ratios at the same value 0.59.

**Supplemental Figure 2.**
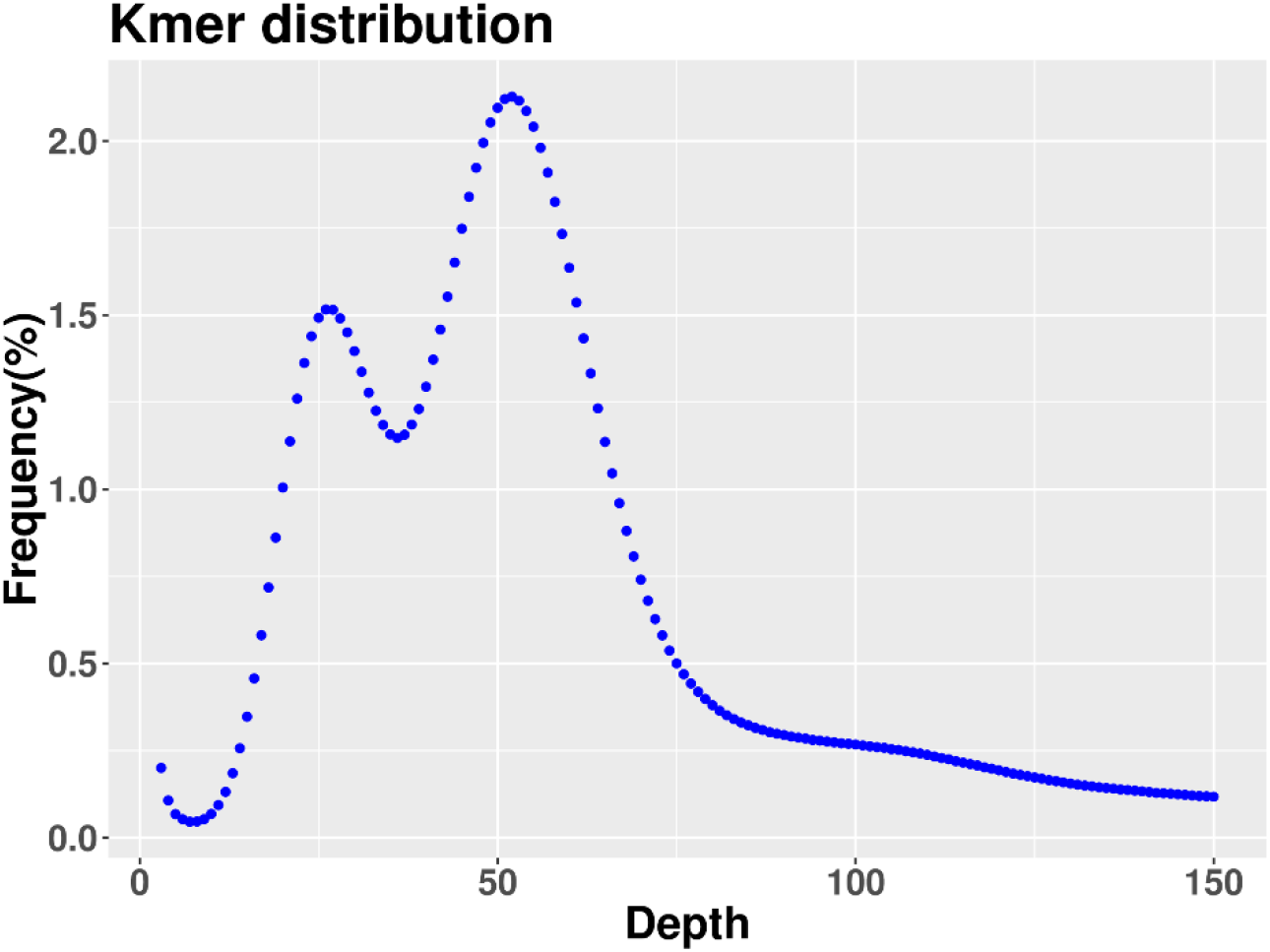
K-mer (k=19) distribution.

**Supplemental Figure 3.**
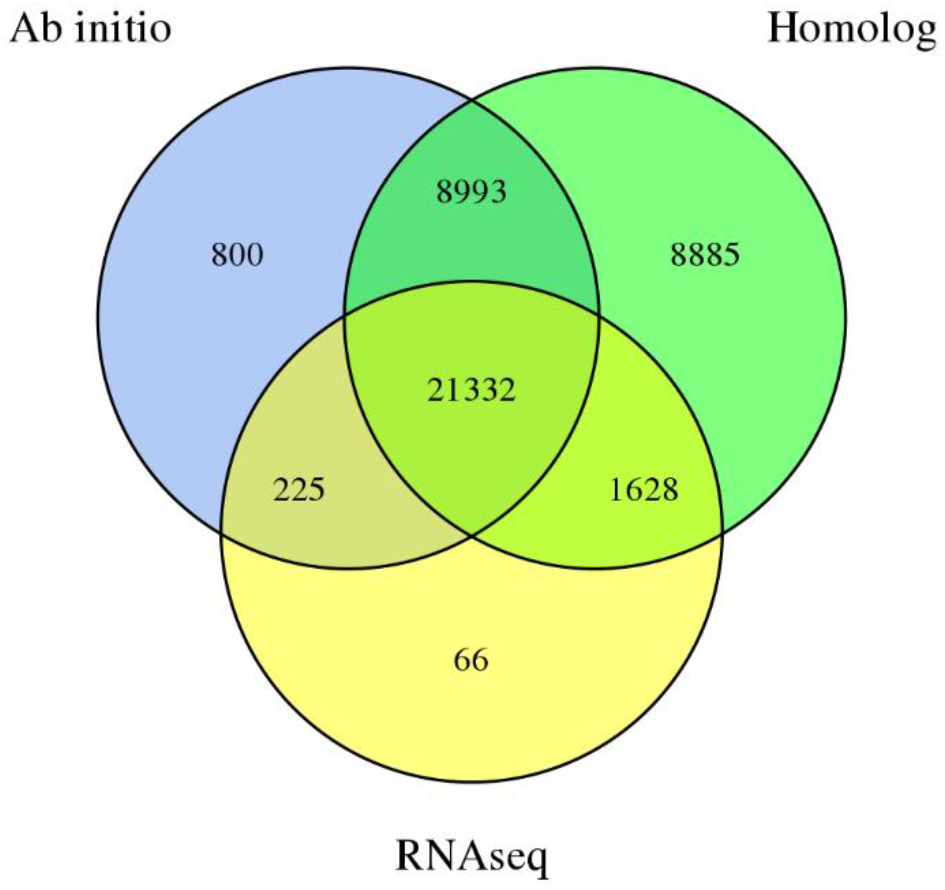
Genes derived from the distribution map of the three prediction methods. A combination of plant protein homology mapping, transcriptome data, and *ab initio* gene prediction was used to generate gene model predictions.

**Supplemental Figure 4.**
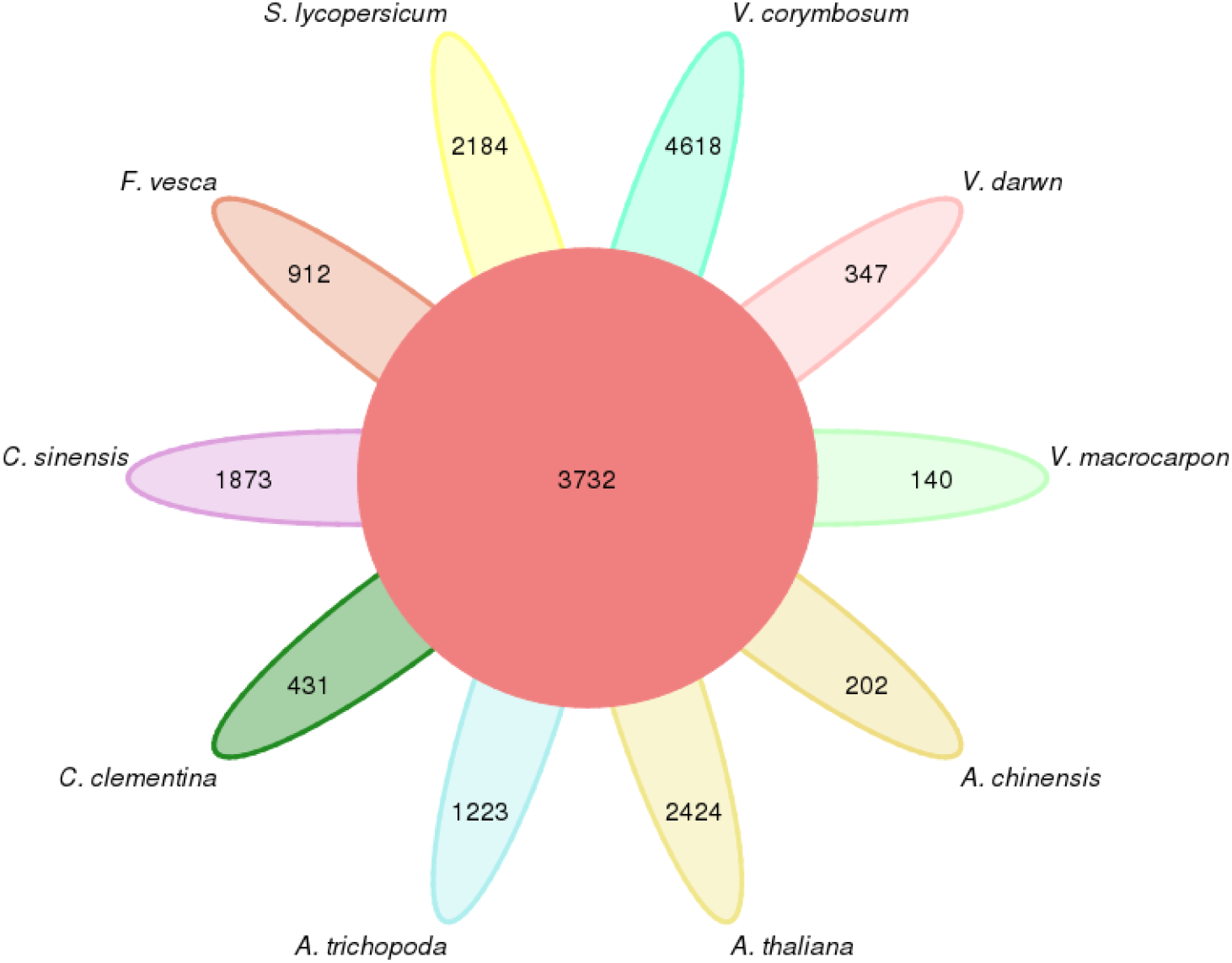
Gene family cluster petal diagram.

**Supplemental Figure 5.**
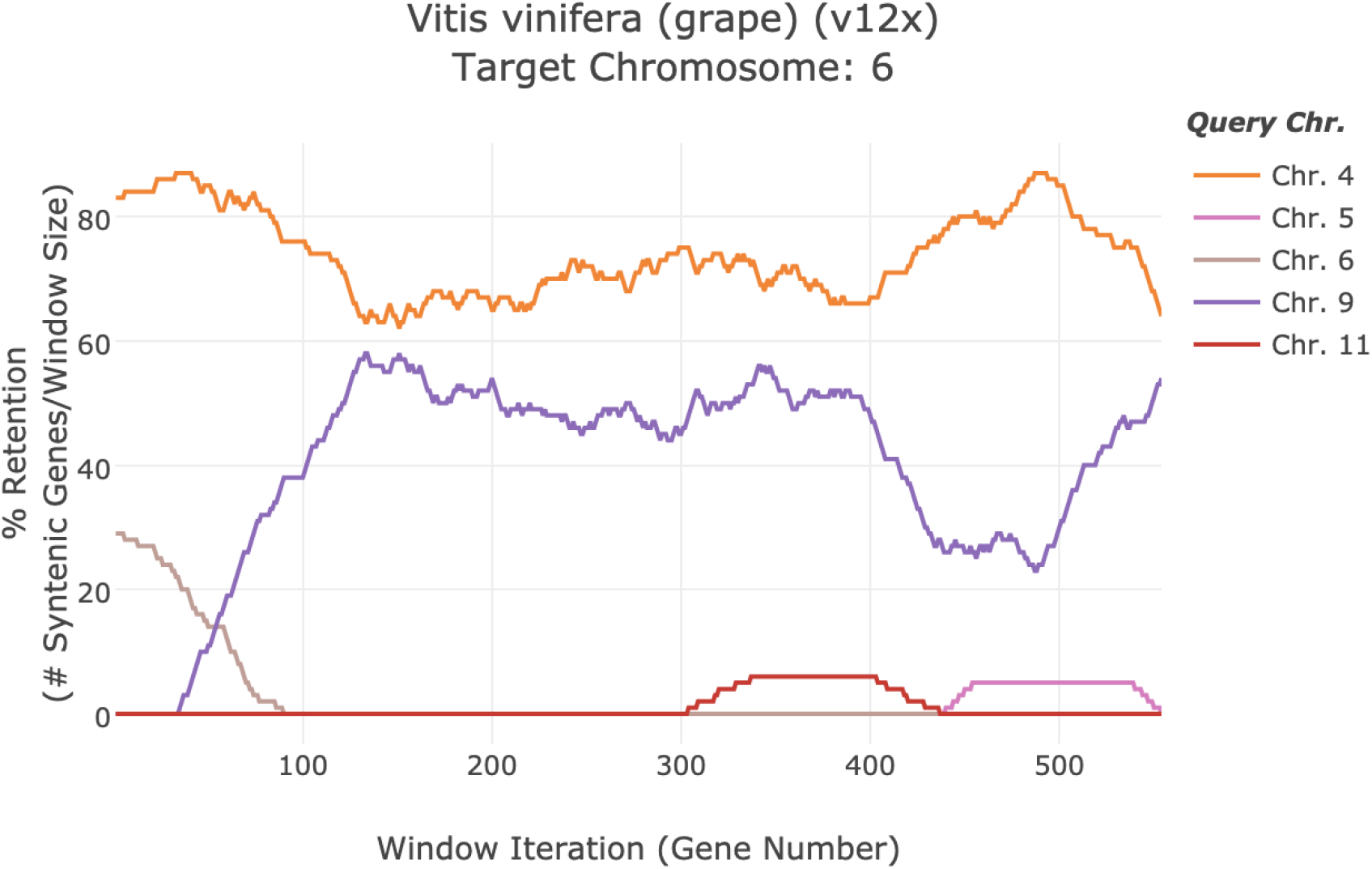
Syntenic comparison of *V. darrowii* against grape (*Vitis vinifera*). The evidence that two chromosomes from *V. darrowii* were syntenic with one grape chromosome, indicates that *V. darrowii* had experienced a new WGD event after the paleohexaploid (γ) event.

## Reference

1 Wu T, Gao Y, Guo X, Zhang M, Gong L. Blackberry and Blueberry Anthocyanin Supplementation Counteract High-Fat-Diet-Induced Obesity by Alleviating Oxidative Stress and Inflammation and Accelerating Energy Expenditure. Oxidative Medicine and Cellular Longevity. 2018; 2018: e4051232.

2 Kalt W, Cassidy A, Howard LR et al. Recent Research on the Health Benefits of Blueberries and Their Anthocyanins. Advances in Nutrition 2020; 11: 224–236.

3 Romo-Muñoz R, Dote-Pardo J, Garrido-Henríquez H, Araneda-Flores J, Gil JM. Blueberry consumption and healthy lifestyles in an emerging market. Span J Agric Res 2020; 17: e0111.

4 Li Y, Sun H, Chen L. The blueberry industry of China: the past 10 years and the future. Acta Hortic 2017; : 531–536.

5 Li Y, Pei J, Chen L, Sun H. China blueberry industry report 2020. Journal of Jilin Agricultural University 2021; 43: 1–8.

6 Strik B. Blueberry production and research trends in north America. Acta Hortic 2006; : 173–184.

7 Retamales JB, Palma MJ, Morales YA, Lobos GA, Moggia CE, Mena CA. Blueberry production in Chile: current status and future developments. Rev Bras Frutic 2014; 36: 58–67.

8 Cho HY, Kadowaki M, Graduate School of Agriculture, Tokyo University of Agriculture and Technology, Fuchu 183-8509, Japan et al. Influence of light quality on flowering characteristics, potential for year-round fruit production and fruit quality of blueberry in a plant factory. Fruits 2019; 74: 3–10.

9 Fang Y, Nunez GH, Silva MN da, Phillips DA, Munoz PR. A review for southern highbush blueberry alternative production systems. Agronomy 2020; 10: 1531.

10 Sharpe RH, Darrow GM. Breeding blueberries for the Florida climate. Proceedings of the Florida State Horticultural Society, 1959 1960; 72: 308–11.

11 Lobos GA, Hancock JF. Breeding blueberries for a changing global environment: a review. Front Plant Sci 2015; 6. doi:10.3389/fpls.2015.00782.

12 Chavez DJ, Lyrene PM. Effects of self-pollination and cross-pollination of Vaccinium darrowii (Ericaceae) and other low-chill blueberries. HortScience 2009; 44: 1538–1541.

13 Sultana N, Pascual-Díaz JP, Gers A et al. Contribution to the knowledge of genome size evolution in edible blueberries (genus Vaccinium). Journal of Berry Research 2019; Preprint: 1–15.

14 Colle M, Leisner CP, Wai CM et al. Haplotype-phased genome and evolution of phytonutrient pathways of tetraploid blueberry. Gigascience 2019; 8. doi:10.1093/gigascience/giz012.

15 Diaz-Garcia L, Garcia-Ortega LF, González-Rodríguez M, Delaye L, Iorizzo M, Zalapa J. Chromosome-level genome assembly of the american cranberry (Vaccinium macrocarpon Ait.) and its wild relative Vaccinium microcarpum. Front Plant Sci 2021; 12: 633310.

16 Wu C, Deng C, Hilario E et al. A chromosome-scale assembly of the bilberry genome identifies a complex locus controlling berry anthocyanin composition. Molecular Ecology Resources 2021; n/a. doi:10.1111/1755-0998.13467.

17 Wilkie JD, Sedgley M, Olesen T. Regulation of floral initiation in horticultural trees. J Exp Bot 2008; 59: 3215–3228.

18 Song G, Walworth A, Zhao D, Jiang N, Hancock JF. The Vaccinium corymbosumFLOWERING LOCUS T-like gene (VcFT): a flowering activator reverses photoperiodic and chilling requirements in blueberry. Plant Cell Rep 2013; 32: 1759–1769.

19 Chavez DJ, Lyrene PM. Interspecific Crosses and Backcrosses between Diploid Vaccinium darrowii and Tetraploid Southern Highbush Blueberry. Journal of the American Society for Horticultural Science 2009; 134: 273–280.

20 Galbraith DW, Harkins KR, Maddox JM, Ayres NM, Sharma DP, Firoozabady E. Rapid Flow Cytometric Analysis of the Cell Cycle in Intact Plant Tissues. Science 1983; 220: 1049–1051.

21 Robinson JP, Grégori G. Principles of Flow Cytometry. In: Flow Cytometry with Plant Cells. John Wiley & Sons, Ltd, 2007, pp 19–40.

22 Porebski S, Bailey LG, Baum BR. Modification of a CTAB DNA extraction protocol for plants containing high polysaccharide and polyphenol components. Plant Mol Biol Rep 1997; 15: 8–15.

23 Berensmeier S. Magnetic particles for the separation and purification of nucleic acids. Appl Microbiol Biotechnol 2006; 73: 495–504.

24 Altschul SF, Gish W, Miller W, Myers EW, Lipman DJ. Basic local alignment search tool. Journal of Molecular Biology 1990; 215: 403–410.

25 Li R, Li Y, Kristiansen K, Wang J. SOAP: short oligonucleotide alignment program. Bioinformatics 2008; 24: 713–714.

26 Deamer D, Akeson M, Branton D. Three decades of nanopore sequencing. Nature Biotechnology 2016; 34: 518–524.

27 Jaakola L, Pirttilä AM, Halonen M, Hohtola A. Isolation of high quality RNA from bilberry (Vaccinium myrtillus L.) fruit. Mol Biotechnol 2001; 19: 201–203.

28 Koren S, Walenz BP, Berlin K, Miller JR, Bergman NH, Phillippy AM. Canu: scalable and accurate long-read assembly via adaptive k-mer weighting and repeat separation. Genome Res 2017; 27: 722–736.

29 Walker BJ, Abeel T, Shea T et al. Pilon: an integrated tool for comprehensive microbial variant detection and genome assembly improvement. PLoS One 2014; 9: e112963.

30 Vaser R, Sović I, Nagarajan N, Šikić M. Fast and accurate de novo genome assembly from long uncorrected reads. Genome Res 2017; 27: 737–746.

31 Parra G, Bradnam K, Korf I. CEGMA: a pipeline to accurately annotate core genes in eukaryotic genomes. Bioinformatics 2007; 23: 1061–1067.

32 Simão FA, Waterhouse RM, Ioannidis P, Kriventseva EV, Zdobnov EM. BUSCO: assessing genome assembly and annotation completeness with single-copy orthologs. Bioinformatics 2015; 31: 3210–3212.

33 Rao SSP, Huntley MH, Durand NC et al. A 3D map of the human genome at kilobase resolution reveals principles of chromatin looping. Cell 2014; 159: 1665–1680.

34 Li H, Durbin R. Fast and accurate short read alignment with Burrows-Wheeler transform. Bioinformatics 2009; 25: 1754–1760.

35 Servant N, Varoquaux N, Lajoie BR et al. HiC-Pro: an optimized and flexible pipeline for Hi-C data processing. Genome Biol 2015; 16: 259.

36 Burton JN, Adey A, Patwardhan RP, Qiu R, Kitzman JO, Shendure J. Chromosome-scale scaffolding of de novo genome assemblies based on chromatin interactions. Nat Biotechnol 2013; 31: 1119–1125.

37 Price AL, Jones NC, Pevzner PA. De novo identification of repeat families in large genomes. Bioinformatics 2005; 21 Suppl 1: i351–358.

38 Xu Z, Wang H. LTR_FINDER: an efficient tool for the prediction of full-length LTR retrotransposons. Nucleic Acids Res 2007; 35: W265–268.

39 Jurka J, Kapitonov VV, Pavlicek A, Klonowski P, Kohany O, Walichiewicz J. Repbase Update, a database of eukaryotic repetitive elements. Cytogenet Genome Res 2005; 110: 462–467.

40 Hoede C, Arnoux S, Moisset M et al. PASTEC: an automatic transposable element classification tool. PLoS One 2014; 9: e91929.

41 Tarailo-Graovac M, Chen N. Using RepeatMasker to identify repetitive elements in genomic sequences. Curr Protoc Bioinformatics 2009; Chapter 4: Unit 4.10.

42 Lowe TM, Eddy SR. tRNAscan-SE: a program for improved detection of transfer RNA genes in genomic sequence. Nucleic Acids Res 1997; 25: 955–964.

43 Griffiths-Jones S, Moxon S, Marshall M, Khanna A, Eddy SR, Bateman A. Rfam: annotating non-coding RNAs in complete genomes. Nucleic Acids Res 2005; 33: D121–124.

44 Birney E, Clamp M, Durbin R. GeneWise and Genomewise. Genome Res 2004; 14: 988–995.

45 She R, Chu JS-C, Wang K, Pei J, Chen N. GenBlastA: enabling BLAST to identify homologous gene sequences. Genome Res 2009; 19: 143–149.

46 Haas BJ, Salzberg SL, Zhu W et al. Automated eukaryotic gene structure annotation using EVidenceModeler and the Program to Assemble Spliced Alignments. Genome Biol 2008; 9: R7.

47 Burge C, Karlin S. Prediction of complete gene structures in human genomic DNA. J Mol Biol 1997; 268: 78–94.

48 Stanke M, Waack S. Gene prediction with a hidden Markov model and a new intron submodel. Bioinformatics 2003; 19 Suppl 2: ii215–225.

49 Majoros WH, Pertea M, Salzberg SL. TigrScan and GlimmerHMM: two open source ab initio eukaryotic gene-finders. Bioinformatics 2004; 20: 2878–2879.

50 Blanco E, Parra G, Guigó R. Using geneid to identify genes. Curr Protoc Bioinformatics 2007; Chapter 4: Unit 4.3.

51 Korf I. Gene finding in novel genomes. BMC Bioinformatics 2004; 5: 59.

52 Keilwagen J, Hartung F, Paulini M, Twardziok SO, Grau J. Combining RNA-seq data and homology-based gene prediction for plants, animals and fungi. BMC Bioinformatics 2018; 19: 189.

53 Kim D, Langmead B, Salzberg SL. HISAT: a fast spliced aligner with low memory requirements. Nat Methods 2015; 12: 357–360.

54 Pertea M, Pertea GM, Antonescu CM, Chang T-C, Mendell JT, Salzberg SL. StringTie enables improved reconstruction of a transcriptome from RNA-seq reads. Nat Biotechnol 2015; 33: 290–295.

55 Tang S, Lomsadze A, Borodovsky M. Identification of protein coding regions in RNA transcripts. Nucleic Acids Res 2015; 43: e78.

56 Campbell MA, Haas BJ, Hamilton JP, Mount SM, Buell CR. Comprehensive analysis of alternative splicing in rice and comparative analyses with Arabidopsis. BMC Genomics 2006; 7: 327.

57 Marchler-Bauer A, Lu S, Anderson JB et al. CDD: a Conserved Domain Database for the functional annotation of proteins. Nucleic Acids Res 2011; 39: D225–229.

58 Koonin EV, Fedorova ND, Jackson JD et al. A comprehensive evolutionary classification of proteins encoded in complete eukaryotic genomes. Genome Biol 2004; 5: R7.

59 Dimmer EC, Huntley RP, Alam-Faruque Y et al. The UniProt-GO Annotation database in 2011. Nucleic Acids Res 2012; 40: D565–570.

60 Ogata H, Goto S, Sato K, Fujibuchi W, Bono H, Kanehisa M. KEGG: Kyoto Encyclopedia of Genes and Genomes. Nucleic Acids Res 1999; 27: 29–34.

61 Boeckmann B, Bairoch A, Apweiler R et al. The SWISS-PROT protein knowledgebase and its supplement TrEMBL in 2003. Nucleic Acids Res 2003; 31: 365–370.

62 Yang Z. PAML 4: phylogenetic analysis by maximum likelihood. Mol Biol Evol 2007; 24: 1586–1591.

63 De Bie T, Cristianini N, Demuth JP, Hahn MW. CAFE: a computational tool for the study of gene family evolution. Bioinformatics 2006; 22: 1269–1271.

64 Wang Y, Tang H, Debarry JD et al. MCScanX: a toolkit for detection and evolutionary analysis of gene synteny and collinearity. Nucleic Acids Res 2012; 40: e49.

65 Klopfenstein DV, Zhang L, Pedersen BS et al. GOATOOLS: A Python library for Gene Ontology analyses. Sci Rep 2018; 8: 10872.

66 Cui F, Brosché M, Lehtonen MT et al. Dissecting Abscisic Acid Signaling Pathways Involved in Cuticle Formation. Mol Plant 2016; 9: 926–938.

67 Hellemans J, Mortier G, De Paepe A, Speleman F, Vandesompele J. qBase relative quantification framework and software for management and automated analysis of real-time quantitative PCR data. Genome Biol 2007; 8: R19.

68 Lyrene PM, Vorsa N, Ballington JR. Polyploidy and sexual polyploidization in the genus *Vaccinium*. Euphytica 2003; 133: 27–36.

69 Vorsa N, Johnson-Cicalese J, Polashock J. A blueberry by cranberry hybrid derived from a Vaccinium darrowii × (V. macrocarpon × V. oxycoccos) intersectional cross. Acta Hortic 2009; : 187–190.

70 Sakhanokho HF, Rinehart TA, Stringer SJ, Islam-Faridi MN, Pounders CT. Variation in nuclear DNA content and chromosome numbers in blueberry. Scientia Horticulturae 2018; 233: 108–113.

71 Kent WJ. BLAT--the BLAST-like alignment tool. Genome Res 2002; 12: 656–664.

72 Bowers JE, Chapman BA, Rong J, Paterson AH. Unravelling angiosperm genome evolution by phylogenetic analysis of chromosomal duplication events. Nature 2003; 422: 433–438.

73 Archetti M, Döring TF, Hagen SB et al. Unravelling the evolution of autumn colours: an interdisciplinary approach. Trends in Ecology & Evolution 2009; 24: 166–173.

74 Rabino I, Mancinelli AL. Light, Temperature, and Anthocyanin Production. Plant Physiology 1986; 81: 922–924.

75 Rowan DD, Cao M, Lin-Wang K et al. Environmental regulation of leaf colour in red 35S:PAP1 Arabidopsis thaliana. New Phytol 2009; 182: 102–115.

76 Lin-Wang K, Micheletti D, Palmer J et al. High temperature reduces apple fruit colour via modulation of the anthocyanin regulatory complex. Plant, Cell & Environment 2011; 34: 1176–1190.

77 Chen M, Thelen JJ. ACYL-LIPID DESATURASE2 Is Required for Chilling and Freezing Tolerance in Arabidopsis[C][W]. Plant Cell 2013; 25: 1430–1444.

78 Román Á, Hernández ML, Soria-García Á et al. Non-redundant Contribution of the Plastidial FAD8 ω-3 Desaturase to Glycerolipid Unsaturation at Different Temperatures in Arabidopsis. Molecular Plant 2015; 8: 1599–1611.

79 Ramakrishna A, Ravishankar GA. Influence of abiotic stress signals on secondary metabolites in plants. Plant Signal Behav 2011; 6: 1720–1731.

80 Aguirre-Becerra H, Vazquez-Hernandez MC, Saenz de la O D et al. Role of Stress and Defense in Plant Secondary Metabolites Production. In: Pal D, Nayak AK (eds). Bioactive Natural Products for Pharmaceutical Applications. Springer International Publishing: Cham, 2021, pp 151–195.

81 Supek F, Bošnjak M, Škunca N, Šmuc T. REVIGO summarizes and visualizes long lists of gene ontology terms. PLoS One 2011; 6: e21800.

82 Han S-H, Park Y-J, Park C-M. HOS1 activates DNA repair systems to enhance plant thermotolerance. Nat Plants 2020; 6: 1439–1446.

83 Nicolau M, Picault N, Descombin J et al. The plant mobile domain proteins MAIN and MAIL1 interact with the phosphatase PP7L to regulate gene expression and silence transposable elements in Arabidopsis thaliana. PLOS Genetics 2020; 16: e1008324.

84 Xu D, Marino G, Klingl A et al. Extrachloroplastic PP7L Functions in Chloroplast Development and Abiotic Stress Tolerance. Plant Physiology 2019; 180: 323–341.

85 Møller SG, Kim Y-S, Kunkel T, Chua N-H. PP7 Is a Positive Regulator of Blue Light Signaling in Arabidopsis. The Plant Cell 2003; 15: 1111–1119.

86 Genoud T, Cruz MTS, Kulisic T, Sparla F, Fankhauser C, Métraux J-P. The Protein Phosphatase 7 Regulates Phytochrome Signaling in Arabidopsis. PLOS ONE 2008; 3: e2699.

87 Ifuku K, Nakatsu T, Shimamoto R et al. Structure and function of the PsbP protein of Photosystem II from higher plants. Photosynth Res 2005; 84: 251–255.

88 Ifuku K, Yamamoto Y, Ono T, Ishihara S, Sato F. PsbP Protein, But Not PsbQ Protein, Is Essential for the Regulation and Stabilization of Photosystem II in Higher Plants. Plant Physiology 2005; 139: 1175–1184.

89 Yamamoto M, Uji S, Sugiyama T et al. ERdj3B-Mediated Quality Control Maintains Anther Development at High Temperatures1 [OPEN]. Plant Physiology 2020; 182: 1979–1990.

90 Taulavuori K, Laine K, Taulavuori E. Experimental studies on Vaccinium myrtillus and Vaccinium vitis-idaea in relation to air pollution and global change at northern high latitudes: A review. Environmental and Experimental Botany 2013; 87: 191–196.

91 Yang Q, Liu E, Fu Y, Yuan F, Zhang T, Peng S. High temperatures during flowering reduce fruit set in rabbiteye blueberry. Journal of the American Society for Horticultural Science 2019; 144: 339–351.

92 Walworth AE, Chai B, Song G. Transcript Profile of Flowering Regulatory Genes in VcFT-Overexpressing Blueberry Plants. PLOS ONE 2016; 11: e0156993.

93 Gao X, Walworth AE, Mackie C, Song G. Overexpression of blueberry FLOWERING LOCUS T is associated with changes in the expression of phytohormone-related genes in blueberry plants. Hortic Res 2016; 3: 1–9.

94 Schomburg FM, Patton DA, Meinke DW, Amasino RM. FPA, a Gene Involved in Floral Induction in Arabidopsis, Encodes a Protein Containing RNA-Recognition Motifs. The Plant Cell 2001; 13: 1427–1436.

95 Veley KM, Michaels SD. Functional Redundancy and New Roles for Genes of the Autonomous Floral-Promotion Pathway. Plant Physiology 2008; 147: 682–695.

96 Sonmez C, Bäurle I, Magusin A et al. RNA 3′ processing functions of Arabidopsis FCA and FPA limit intergenic transcription. PNAS 2011; 108: 8508–8513.

97 Wu Z, Fang X, Zhu D, Dean C. Autonomous Pathway: FLOWERING LOCUS C Repression through an Antisense-Mediated Chromatin-Silencing Mechanism1 [CC-BY]. Plant Physiology 2020; 182: 27–37.

98 Yoo SK, Chung KS, Kim J et al. CONSTANS activates SUPPRESSOR OF OVEREXPRESSION OF CONSTANS 1 through FLOWERING LOCUS T to promote flowering in Arabidopsis. Plant Physiology 2005; 139: 770–778.

99 Hou D, Li L, Ma T et al. The SOC1-like gene BoMADS50 is associated with the flowering of Bambusa oldhamii. Hortic Res 2021; 8: 1–13.

100 Hepworth SR, Valverde F, Ravenscroft D, Mouradov A, Coupland G. Antagonistic regulation of flowering-time gene SOC1 by CONSTANS and FLC via separate promoter motifs. The EMBO journal 2002; 21. doi:10.1093/emboj/cdf432.

101 Immink RGH, Posé D, Ferrario S et al. Characterization of SOC1’s central role in flowering by the identification of its upstream and downstream regulators. Plant Physiology 2012; 160: 433–449.

102 Horvath D. Common mechanisms regulate flowering and dormancy. Plant Science 2009; 177: 523–531.

103 Trainin T, Bar-Ya’akov I, Holland D. ParSOC1, a MADS-box gene closely related to Arabidopsis AGL20/SOC1, is expressed in apricot leaves in a diurnal manner and is linked with chilling requirements for dormancy break. Tree Genetics & Genomes 2013; 9: 753–766.

104 Chen M-K, Hsu W-H, Lee P-F, Thiruvengadam M, Chen H-I, Yang C-H. The MADS box gene, FOREVER YOUNG FLOWER, acts as a repressor controlling floral organ senescence and abscission in Arabidopsis. Plant J 2011; 68: 168–185.

105 Suter L, Rüegg M, Zemp N, Hennig L, Widmer A. Gene regulatory variation mediates flowering responses to vernalization along an altitudinal gradient in Arabidopsis. Plant Physiol 2014; 166: 1928–1942.

106 Voogd C, Wang T, Varkonyi-Gasic E. Functional and expression analyses of kiwifruit SOC1-like genes suggest that they may not have a role in the transition to flowering but may affect the duration of dormancy. Journal of Experimental Botany 2015; 66: 4699–4710.

107 Horvath DP, Anderson JV, Chao WS, Foley ME. Knowing when to grow: signals regulating bud dormancy. Trends in Plant Science 2003; 8: 534–540.

108 Gregis V, Andrés F, Sessa A et al. Identification of pathways directly regulated by SHORT VEGETATIVE PHASE during vegetative and reproductive development in Arabidopsis. Genome Biol 2013; 14: R56.

109 Wang SY, Camp MJ, Ehlenfeldt MK. Antioxidant capacity and α-glucosidase inhibitory activity in peel and flesh of blueberry (Vaccinium spp.) cultivars. Food Chemistry 2012; 132: 1759–1768.

110 Yeats TH, Rose JKC. The formation and function of plant cuticles. Plant Physiol 2013; 163: 5–20.

111 Jaakola L, Määttä K, Pirttilä AM, Törrönen R, Kärenlampi S, Hohtola A. Expression of genes involved in anthocyanin biosynthesis in relation to anthocyanin, proanthocyanidin, and flavonol levels during bilberry fruit development. Plant Physiol 2002; 130: 729–739.

112 Zhang Y, Butelli E, Martin C. Engineering anthocyanin biosynthesis in plants. Current Opinion in Plant Biology 2014; 19: 81–90.

113 Forlani S, Masiero S, Mizzotti C. Fruit ripening: the role of hormones, cell wall modifications, and their relationship with pathogens. Journal of Experimental Botany 2019; 70: 2993–3006.

114 Gupta V, Estrada AD, Blakley I et al. RNA-Seq analysis and annotation of a draft blueberry genome assembly identifies candidate genes involved in fruit ripening, biosynthesis of bioactive compounds, and stage-specific alternative splicing. GigaScience 2015; 4. doi:10.1186/s13742-015-0046-9.

115 Sapers GM, Phillips JG. Leakage of anthocyanins from skin of raw and cooked highbush blueberries (Vaccinium corymbosum L.). Journal of Food Science 1985; 50: 437–439.

116 Trivedi P, Karppinen K, Klavins L et al. Compositional and morphological analyses of wax in northern wild berry species. Food Chemistry 2019; 295: 441–448.

117 Li R, Sun S, Wang H et al. FIS1 encodes a GA2-oxidase that regulates fruit firmness in tomato. Nature Communications 2020; 11: 5844.

118 Bourdenx B, Bernard A, Domergue F et al. Overexpression of Arabidopsis *ECERIFERUM1* promotes wax very-long-chain alkane biosynthesis and influences plant response to biotic and abiotic stresses. Plant Physiol 2011; 156: 29–45.

119 Bernard A, Domergue F, Pascal S et al. Reconstitution of plant alkane biosynthesis in yeast demonstrates that *Arabidopsis* ECERIFERUM1 and ECERIFERUM3 are core components of a very-long-chain alkane synthesis complex. Plant Cell 2012; 24: 3106–3118.

120 Greer S, Wen M, Bird D et al. The cytochrome P450 enzyme CYP96A15 Is the midchain alkane hydroxylase responsible for formation of secondary alcohols and ketones in stem cuticular wax of Arabidopsis. Plant Physiol 2007; 145: 653–667.

121 Li F, Wu X, Lam P et al. Identification of the Wax Ester Synthase/Acyl-Coenzyme A:Diacylglycerol Acyltransferase WSD1 required for stem wax ester biosynthesis in arabidopsis. Plant Physiol 2008; 148: 97–107.

122 Yeats TH, Huang W, Chatterjee S et al. Tomato Cutin Deficient 1 (CD1) and putative orthologs comprise an ancient family of cutin synthase-like (CUS) proteins that are conserved among land plants. The Plant Journal 2014; 77: 667–675.

123 Hanada K, Zou C, Lehti-Shiu MD, Shinozaki K, Shiu S-H. Importance of lineage-specific expansion of plant tandem duplicates in the adaptive response to environmental stimuli. Plant Physiol 2008; 148: 993–1003.

124 Wang X, Gao Y, Wu X et al. High-quality evergreen azalea genome reveals tandem duplication-facilitated low-altitude adaptability and floral scent evolution. Plant Biotechnology Journal 2021; n/a. doi:10.1111/pbi.13680.

125 Freeling M. Bias in plant gene content following different sorts of duplication: tandem, whole-genome, segmental, or by transposition. Annual review of plant biology 2009; 60: 433–453.

126 Rody HV, Baute GJ, Rieseberg LH, Oliveira LO. Both mechanism and age of duplications contribute to biased gene retention patterns in plants. BMC genomics 2017; 18: 1–10.

127 Kitamura Y, Takeuchi T, Yamane H, Tao R. Simultaneous down-regulation of DORMANCY-ASSOCIATED MADS-box6 and SOC1 during dormancy release in Japanese apricot (Prunus mume) flower buds. The Journal of Horticultural Science and Biotechnology 2016; 91: 476–482.

128 Wang J, Gao Z, Li H et al. Dormancy-associated MADS-box (DAM) genes influence chilling requirement of sweet cherries and co-regulate flower development with SOC1 gene. International Journal of Molecular Sciences 2020; 21: 921.

129 Dorca-Fornell C, Gregis V, Grandi V, Coupland G, Colombo L, Kater MM. The Arabidopsis SOC1-like genes AGL42, AGL71 and AGL72 promote flowering in the shoot apical and axillary meristems. Plant J 2011; 67: 1006–1017.

130 Mouhu K, Kurokura T, Koskela EA, Albert VA, Elomaa P, Hytönen T. The Fragaria vesca homolog of SUPPRESSOR OF OVEREXPRESSION OF CONSTANS1 represses flowering and promotes vegetative growth. The Plant Cell 2013; 25: 3296–3310.

131 Melzer S, Lens F, Gennen J, Vanneste S, Rohde A, Beeckman T. Flowering-time genes modulate meristem determinacy and growth form in Arabidopsis thaliana. Nat Genet 2008; 40: 1489–1492.

132 Chen J, Zhu X, Ren J et al. Suppressor of Overexpression of CO 1 Negatively Regulates Dark-Induced Leaf Degreening and Senescence by Directly Repressing Pheophytinase and Other Senescence-Associated Genes in Arabidopsis. Plant Physiol 2017; 173: 1881–1891.

133 Tyagi S, Sri T, Singh A et al. SUPPRESSOR OF OVEREXPRESSION OF CONSTANS1 influences flowering time, lateral branching, oil quality, and seed yield in Brassica juncea cv. Varuna. Funct Integr Genomics 2019; 19: 43–60.

134 Borner R, Kampmann G, Chandler J et al. A MADS domain gene involved in the transition to flowering in Arabidopsis. The Plant Journal 2000; 24: 591–599.

135 Shen L, Kang YGG, Liu L, Yu H. The J-Domain Protein J3 Mediates the Integration of Flowering Signals in Arabidopsis. The Plant Cell 2011; 23: 499–514.

136 Szakiel A, Pączkowski C, Pensec F, Bertsch C. Fruit cuticular waxes as a source of biologically active triterpenoids. Phytochem Rev 2012; 11: 263–284.

137 Chu W, Gao H, Cao S, Fang X, Chen H, Xiao S. Composition and morphology of cuticular wax in blueberry (Vaccinium spp.) fruits. Food Chemistry 2017; 219: 436–442.

138 Chu W, Gao H, Chen H, Wu W, Fang X. Changes in cuticular wax composition of two blueberry cultivars during fruit ripening and postharvest cold storage. J Agric Food Chem 2018; 66: 2870–2876.

139 Oosumi T, Gruszewski HA, Blischak LA et al. High-efficiency transformation of the diploid strawberry (Fragaria vesca) for functional genomics. Planta 2006; 223: 1219–1230.

140 Shulaev V, Sargent DJ, Crowhurst RN et al. The genome of woodland strawberry ( Fragaria vesca ). Nature Genetics 2011; 43: 109–116.

141 Li Y, Pi M, Gao Q, Liu Z, Kang C. Updated annotation of the wild strawberry Fragaria vesca V4 genome. Horticulture Research 2019; 6: 1–9.

142 Longman KA, Wareing PF. Early Induction of Flowering in Birch Seedlings. Nature 1959; 184: 2037–2038.

143 Salojärvi J, Smolander O-P, Nieminen K et al. Genome sequencing and population genomic analyses provide insights into the adaptive landscape of silver birch. Nature Genetics 2017; 49: 904–912.

144 Zhu C, Zheng X, Huang Y et al. Genome sequencing and CRISPR/Cas9 gene editing of an early flowering Mini-Citrus (Fortunella hindsii). Plant Biotechnology Journal 2019; 17: 2199–2210.

145 Bradshaw HD, Ceulemans R, Davis J, Stettler R. Emerging Model Systems in Plant Biology: Poplar(Populus) as A Model Forest Tree. J Plant Growth Regul 2000; 19: 306–313.

146 Taylor G. Populus: Arabidopsis for Forestry. Do We Need a Model Tree? Annals of Botany 2002; 90: 681–689.

